# Bottom-heaviness stabilizes pelagic fish eggs and generates advective transport at their surface

**DOI:** 10.64898/2026.11.01.763416

**Authors:** Maya Van Gelder, Uri Shavit, Dan Givoli, Daniel Rabinovich, Roi Holzman

## Abstract

Pelagic fish eggs are small spheres (∼ 1*mm*) that drift in the open ocean, carried by large-scale currents. Embedded in these currents are egg-scale turbulent eddies that impose local rotational forcing on the eggs. Accordingly, eggs are considered passive tracer particles that move with these eddies, implying that mass transfer of oxygen and waste are controlled only by molecular diffusion. However, eggs are bottom heavy, characterized by a dense embryo and a buoyant oil globule at opposing poles. We hypothesized that such internal structure generates a gravitational restoring torque, stabilizing the egg and generating relative egg-water motion. To test this hypothesis we exposed fish eggs to controlled flows with shear levels typical of upper mixed-layer and estuarine environments, and analyzed their dynamics. Eggs displayed one of two repeatable behaviors: (i) steady orientation, with a near-constant egg angle, and (ii) revolving motion, with egg rotation at time-varying angular speed. Both behaviors generated relative motion between the eggs and the surrounding water. A rigid-body rotational model balancing external flow-induced torque and gravitational restoring torque showed that transitions between these states depend on local flow fields and developmental stage, with older eggs shifting to revolving behavior at lower external fluid torque. Together, these findings suggest that advective transport is important and that diffusion-only models underestimate exchange rates, which are a major component in predicting the effects of environmental factors on embryo survival.

---

The vast majority of marine fish reproduce by external fertilization of eggs that drift in the open water (∼75% [Miller and Kendall, 2009, p. 19]). Typically, the broadcasted eggs are spherical and small (median size = 0.95 mm, with *>* 70% smaller than 1.5 mm; Ahlstrom and Moser [1980], Motani and Wainwright [2015]), and lack parental care Miller and Kendall [2009], Blaxter [2023]. During the egg stage, the embryo undergoes rapid, temperature-dependent development, with typical times of ∼ 50 hours in tropical and temperate climates Pauly and Pullin [1988]. The embryo lacks functional respiratory or circulatory systems, meaning that oxygen acquisition and metabolic waste removal depend on transport across the egg surface. This constraint makes exchange with the surrounding water a key determinant of embryonic performance and survival. Numerous studies have shown that oxygen availability limits embryo development and is directly related to egg survivorship Daykin [1965], Kaur and Toor [1978], Wieland et al. [1994], Hassell et al. [2008], Torrans and Steeby [2008], Martin et al. [2020]. Simultaneously, faster growth and development, resulting from faster metabolism, are also directly related to survivorship of the embryo and larvae (Ref.). The rate at which oxygen can be supplied (and waste removed) is therefore central to understanding early-life history bottlenecks, recruitment variability, and how environmental change may alter reproductive success in fish.

The transfer of solutes (such as oxygen and ammonia) between the egg and the surrounding water is typically modeled as diffusion and advection across a boundary layer at the egg–water interface, with the embryo acting as a sink that maintains the concentration gradient Motani and Wainwright [2015], Kranenbarg et al. [2000].

Current treatments of oxygen transport to pelagic eggs implicitly treat the eggs as passive tracer particles that move with the surrounding water, such that they accelerate and rotate at the same rates as the fluid immediately adjacent to them Motani and Wainwright [2015], Kranenbarg et al. [2000]. Under this assumption, no relative motion between the egg and the water occur, and advection is negligible. Thus, diffusion has been the only mechanism considered to govern mass transfer. The critical question for pelagic eggs is whether the water is effectively “stagnant” relative to the egg surface or whether relative motion exists at the egg’s length scale. This is important because when water flows past a surface, advection can reduce the boundary-layer thickness and increase the concentration gradient at the interface, increasing the flux.

In the ocean, water motion is present across spatial and temporal scales. Pelagic eggs are transported and distributed by larger-scale currents, waves and eddies. At the 1 mm length scale of the egg, the local flow field is dominated by turbulent eddies that scale between 1 to 10 mm at the conditions typical of the upper mixed layer and the epipelagic zone Dillon and Caldwell [1980], Thorpe [2007], where most fish eggs are found. These eddies generate local rotational forcing, potentially creating a rotational motion of the egg around its center. Recent work in particle–turbulence interactions shows that finite-size particles can exhibit spinning and tumbling relative to the surrounding flow, suggesting that egg’s passive tracer behavior is not guaranteed Durham et al. [2009], Van Hout et al. [2022], Byron et al. [2015].

While fish eggs are generally neutrally buoyant, they are often characterized by a non-uniform internal density distribution. The vast majority (∼ 75%) of marine pelagic eggs contain at least one oil globule Ahlstrom and Moser [1980], which is less dense than seawater, while the embryonic tissues, which become spatially localized during development, are denser than seawater. This internal asymmetry, with the oil globule at one pole and the denser embryo at the opposite pole (Fig. 1, 2), can shift the egg’s center of mass away from its geometric center. Such an offset is commonly referred to as bottom-heaviness, describing a body whose center of mass is displaced relative to its center of buoyancy, producing a gravitational restoring torque when the body is tilted away from its stable orientation Kessler [1985], Pedley and Kessler [1992], Durham et al. [2009]. In pelagic fish eggs, this restoring torque is expected to rotate the egg back toward a stable configuration with the oil globule upward. When external flow-induced torque acts on the egg, for example under sheer flow, the balance between the external torque and this gravitational restoring torque could generate relative rotational motion between the water and the egg. Such motion is expected to reduce the thickness of the diffusive boundary layer and give rise to convective transfer, thereby enhancing oxygen flux to the developing embryo.

**Figure 1.**
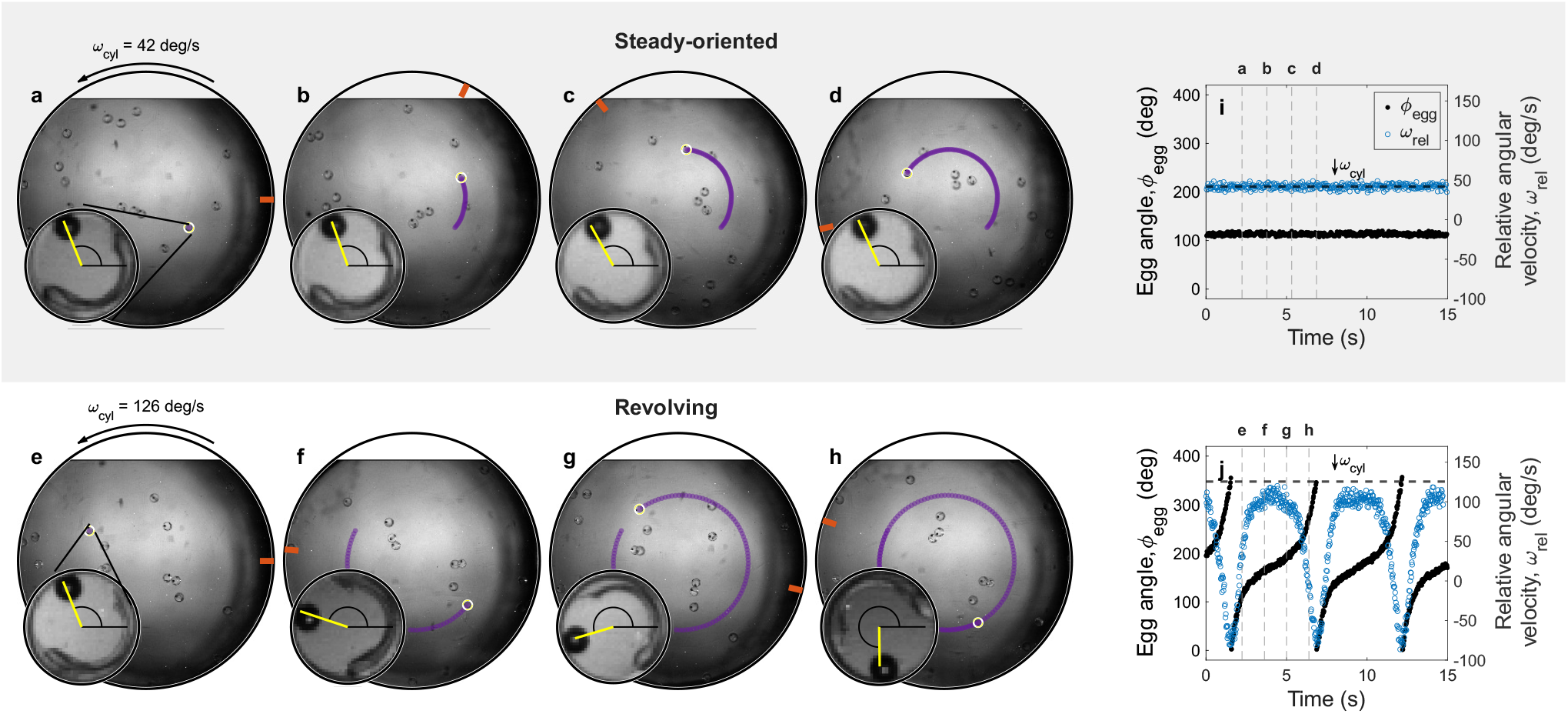
Kinematics of fish egg rotation within a rotating cylinder. (a–d) Time-series snapshots of a stable egg. The outer circle is the cylinder boundary with the orange/red needle representing the instantaneous orientation of the rotating cylinder. The egg that being tracked is marked by a small yellow circle and presented in zoom-in in the smaller circle. The yellow line connects the center of the egg and the center of the oil globule, with the arc representing the orientation of the egg. It is visible that both centers were not necessary accurately recognized by the algorithm, which is the source of the noise represented in (e) the egg kinematics. The angular orientation *ϕ*_*egg*_ of the egg (black) throughout 2.5 cycles of the cylinder, the angular velocity of the water relative to the egg *ω*_rel_ (blue) and the angular velocity of the cylinder *ω*_*cyl*_ (dashed blue line). (f–i) Snapshots of a rotating egg. (j) Corresponding kinematics for the rotating case throughout 5.2 cycles of the cylinder. The magnitude of *ω*_rel_ reached a minimum when the egg was vertically oriented, at *ϕ*_egg_ = 90 *deg* or 270 *deg*, and a maximum when the egg was horizontally oriented, at *ϕ*_egg_ = 0 *deg* or 180 *deg*. Given a counter-clockwise cylinder rotation, the direction of the flow, relative to the egg, was positive when the egg angle was 180 deg and negative when the egg angle was 0 deg. The snapshots values appear in the plot in the gray dashed lines.

**Fig. 2.**
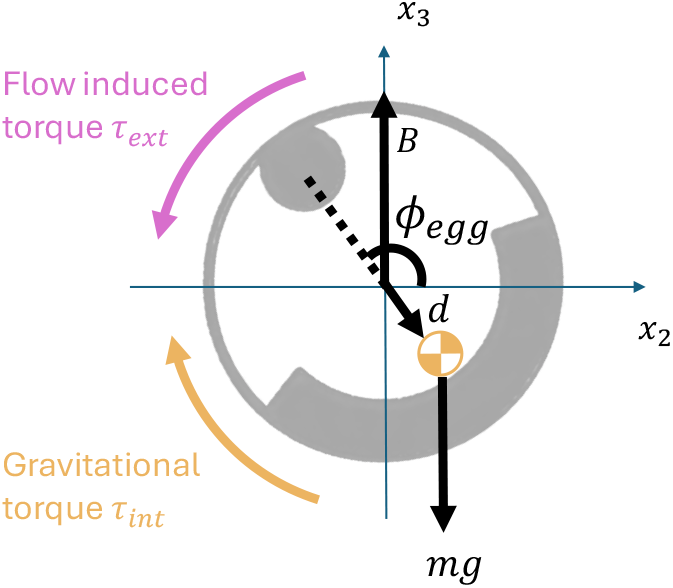
Schematic illustration of the torques applied on a bottom-heavy egg Kessler [1985], Pedley and Kessler [1992], Durham et al. [2009]. The gravitational force (mg) applied at the center of mass (quartered circle) with the offset (d) between the geometric center of the egg and the center of mass generates gravitational restoring torque (*τ*_*int*_) that is dependent on the egg orientation (*ϕ*_*egg*_). Egg orientation is defined by the angle of the oil globule relative to *x*_2_ axis. *τ*_*ext*_ is the external flow-induced torque.

The primary objective of this study is to challenge the prevailing assumption that the egg moves as a passive tracer particle, with no relative motion between the egg and the surrounding water.

To test this hypothesis, we measured the orientation of *S. aurata* fish eggs in a horizontal cylinder rotating at a constant angular velocity, in which the water rotated as a solid body (see *Methods*). The range of angular velocities in the cylinder corresponded to turbulent shear values typically observed in the upper mixed layer of the ocean and in estuaries Dillon and Caldwell [1980], Song and Rau [2022]. Egg orientation was tracked by identifying the angle between the oil globule and the center of the egg (*ϕ*_*egg*_; Fig. 2). The trajectories of all analyzed eggs showed that each egg’s center moved at the same speed as the cylinder while maintaining a constant distance from the cylinder’s center.

We therefore focused on tracking the orientation of the egg at a range of cylinder velocities. Specifically, we asked: (1) Does bottom-heaviness enable eggs to maintain a stable orientation in a rotating cylinder? (2) Does this stabilization generate relative rotational motion between pelagic fish eggs and the surrounding water? and (3) Does the angular velocity of the cylinder and the developmental stage of the egg affect the egg’s stability and speed of relative rotation?

## Results

### Eggs maintained a stable orientation at low and intermediate cylinder angular velocities

For example, an egg measured at a cylinder angular velocity of 42 *deg s*^−1^ maintained a stable angle of *ϕ*_*egg*_ = 113 *±* 2.9 *deg*, irrespective of its position in the cylinder (Fig. 1a-e). By comparison, a passive tracer particle is expected to rotate with the water, causing its orientation to change continuously at the cylinder angular velocity. While maintaining a stable orientation, the mean angle of 1 DPF (day post fertilization) eggs increased with increasing cylinder velocity (Fig. 3C). At the lowest cylinder velocity of 32 *deg s*^−1^, the mean egg angle was 101.3*±*6.0 *deg* (n=30 eggs), peaking to 148*±*14.0 *deg* (n=30 eggs) at a cylinder velocity of 116.3 *deg s*^−1^. In all these cases, the low variance in egg angle within individual tracks (Fig. 1e) indicated that eggs maintained their orientation over time. Thus, those eggs were defined as steady-oriented eggs.

**Figure 3.**
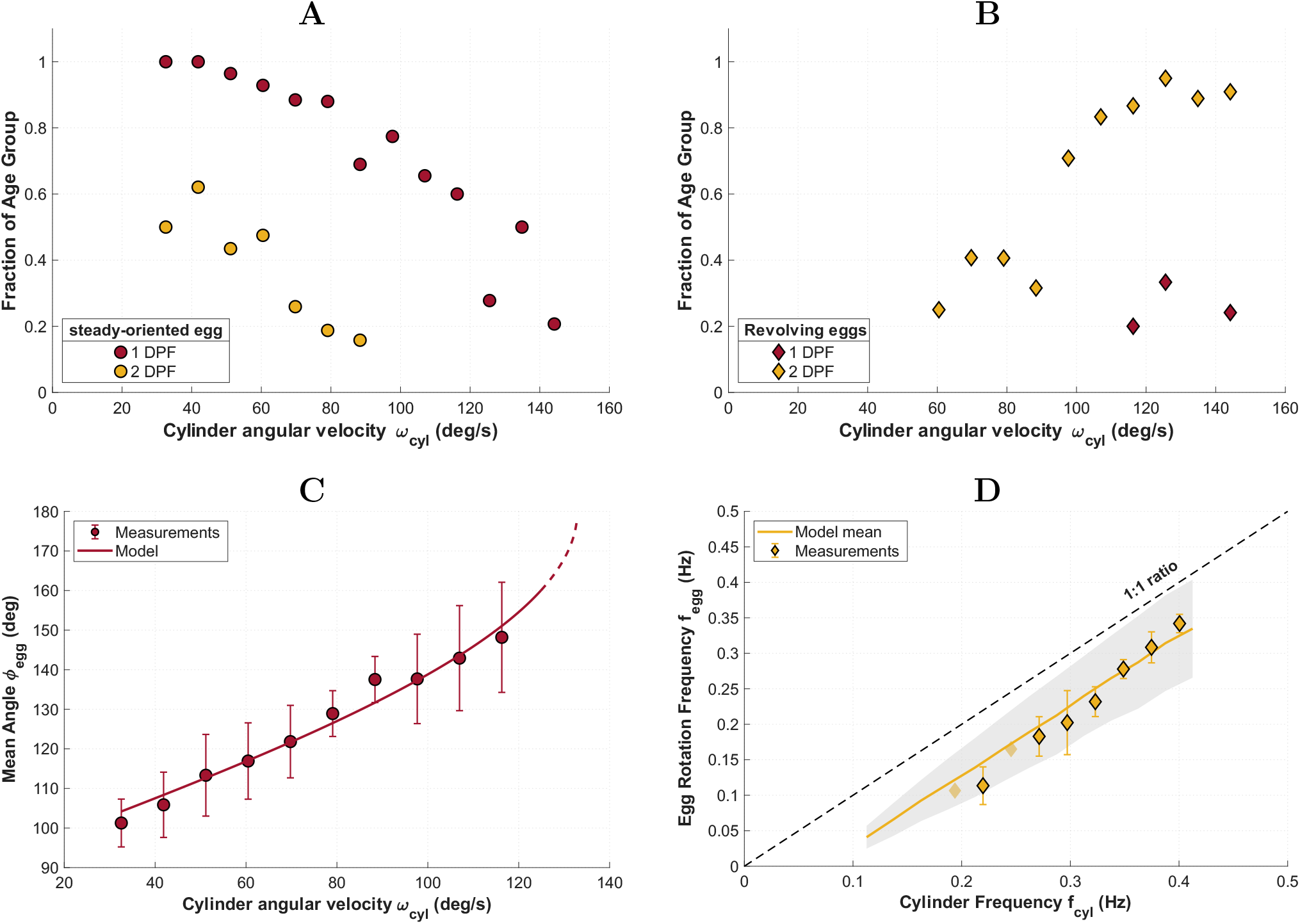
The fraction of eggs in each age group classified as (**A**) steady-oriented eggs and (**B**) revolving eggs as a function of cylinder angular velocity. Fractions in **A** and **B** were calculated from all recorded eggs, including eggs that rotated about the *x*_2_ or *x*_3_ axes and were excluded from the subsequent two-dimensional model comparison, as described in the Results. Eggs are grouped by developmental age: 1 day post fertilization (1 DPF, red) and 2 days post fertilization (2 DPF, yellow). A minority of the eggs (26%) were observed to rotate around the out-of-plane axes of the cylinder, in addition to the primary rotation around the main cylinder axis (see section @@ in the SI). Since our measurements were performed in 2D, such multi-axis rotation could not be accurately resolved and quantified, and these tracks were excluded from further analysis. (**C**) Steady-oriented eggs mean angle, *ϕ*_*egg*_, as a function of cylinder angular velocity, *ω*_*cyl*_ (dots). Only measurement with more than eight eggs per velocity are presented here. The solid line shows the best-fitting solution of Eq. **??**, obtained by jointly fitting *α, a*_*emb*_ and *ρ*_*emb*_ within the parameter range reported in table S?. The best-fitting parameter combination was *α* = 180 deg, *a*_*emb*_ = 0.07 mm and *ρ*_*emb*_ = 1.13 mgmm^−3^ (see SI), yielding *R*^2^ = 0.97. (**D**) Egg rotation frequency as a function of cylinder rotation frequency. Diamonds show the measured mean egg rotation frequency, and error bars indicate one standard deviation. Diamonds without black border represent measurements based on fewer than three eggs. The dashed line marks the 1:1 relation. The yellow line shows the mean model prediction from 100,000 simulations of Eq. **??**, of which 57,456 produced revolving eggs; the gray band shows ±1 standard deviation across the simulations. The model agrees well with the measurements (*R*^2^ = 0.95) and shows a small positive bias, with model - measurement = +0.009 Hz. The parameter ranges used in the simulations are summarized in Table S@.

Because the water rotates with the cylinder, an egg that remains steady in the earthbound frame experiences water moving around it. To quantify this relative motion, we defined the angular velocity of the water relative to the egg as *ω*_rel_ (hereafter relative angular velocity):

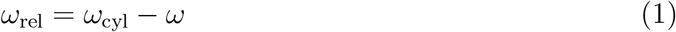

where *ω*_*cyl*_ is the angular velocity of the cylinder and *ω* is the measured angular velocity of the egg, both in the earthbound frame of reference. Thus, a nonzero *ω*_rel_ indicates rotational motion between the water and the egg surface.

For steady-oriented eggs *ω* = 0, and therefore *ω*_rel_ was equal to the cylinder velocity. Correspondingly, the mean angular velocity of the water relative to 1 DPF eggs increased linearly with increasing cylinder velocity from 32 *deg s*^−1^, to 116.3 *deg s*^−1^.

To interpret how the flow conditions determine the observed egg behavior we used the model described in Box. 1. This treatment is similar to that of Kessler [1985], Pedley and Kessler [1992], Durham et al. [2009] for bottom-heavy organisms. Within this framework, steady-orientated eggs are cases in which the flow-induced torque (Eq. 3) is balanced by the gravitational restoring torque generated by bottom-heaviness (Eq. 4); (Fig. 2). While the flow-induced torque is dependent on flow conditions, the gravitational torque is dependent on the offset between the center of mass and center of buoyancy, **d**, and the egg orientation, *ϕ*_*egg*_; Fig. 2. In this case *ω*_rel_ is constant and the left hand side of Eq. 5 is zero. As the external torque increases, the equilibrium angle shifts to larger values, producing greater restoring torque. The solution of Eq. 5 predicted the observed mean angle of stable-oriented 1 DPF eggs well (Fig. 3C). The estimated regression between observed and expected angles had a slope of 0.95 and an of *R*^2^ = 0.97, indicating low model bias.

### Eggs revolved around their center at high cylinder angular velocities

As the cylinder velocity increased, the fraction of steady-oriented eggs decreased, whereas the fraction of eggs revolving around their center increased (Fig. 3A and B). Furthermore, the mean frequency of egg revolution increased with increasing cylinder frequency, but was always lower than the cylinder frequency (Fig. 3D). For example, at the a cylinder angular velocity of 79 *deg s*^−1^ (equivalent to cylinder frequency of 0.22 Hz) the mean frequency of the eggs was 0.11 Hz, and increased to 0.34 Hz at a cylinder angular velocity of 144 *deg s*^−1^ (0.40 Hz).

An example of a revolving egg is shown in Fig. 1e-h. In this case, the egg angle is changing periodically, accelerating and decelerating interchangeably. As a result, the relative angular velocity, *ω*_*rel*_, of the revolving eggs (Eq. 1) varied periodically over time (Fig. 1j). By comparison, a passive tracer particle would be expected to rotate with the cylinder with *ω*_*rel*_ = 0. Because revolving eggs did not rotate at a constant rate, their motion can be quantified by the frequency and amplitude of *ω*_rel_. Unlike the eggs’ frequency, the amplitude of *ω*_*rel*_ (defined as half the difference between its maximum and minimum values over one cycle) was independent of the cylinder velocity (Fig. 4A). Across the range of cylinder velocities, the mean amplitude of *ω*_*rel*_ of 2 DPF eggs was 60.7 *±* 12.7 *deg s*^−1^ (range *@*@ − @@ *deg s*^−1^).

**Figure 4.**
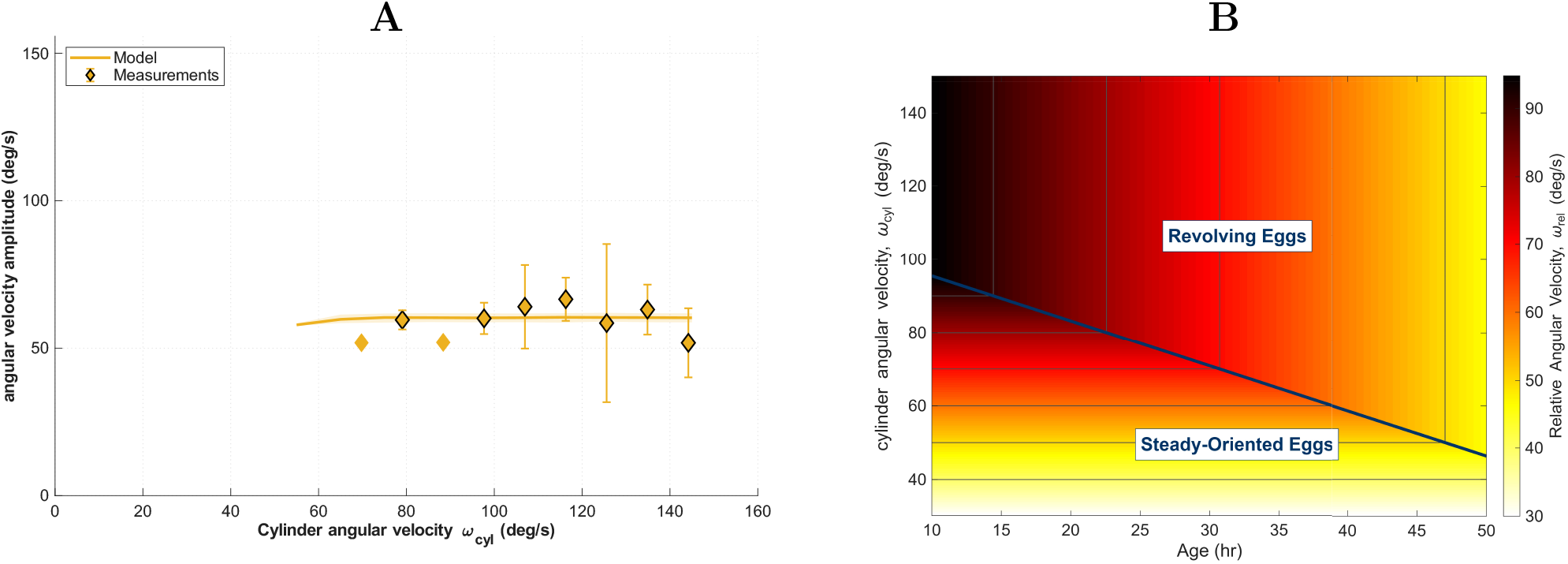
(**A**) Revolving egg angular velocity amplitude as a function of cylinder angular velocity, calculated from Eq. 5. For the relation between age and *d* see SI. Diamonds shows the measured mean amplitude, and error bars indicate one standard deviation. Diamonds without black border represent measurements based on fewer than three eggs. The yellow line shows the mean model prediction from 100,000 simulations of Eq. 5, of which 57,456 produced revolving eggs. The revolving simulations were divided into 10 groups according to their *d* values, and the group shown here is the one that best fits the measurements, excluding measurements based on fewer than three eggs. This group corresponds to *d* = 6.5 *×* 10^−4^– 7.2 *×* 10^−4^ mm and predicts the measurements with an RMSE of 7.3% of the mean measured amplitude. (**B**) The relative rotation magnitude as a function of age and flow conditions as a solution for Eq. 5. The bold blue line divide the two behaviors - under the line all eggs exhibit steady orientation behavior and above it they all exhibit revolving behavior.

To interpret how the flow conditions determines the eggs’ rotation frequency and amplitude, we again used the model described in Box. 1. The model predicts that steady orientation is possible only when the gravitational restoring torque is sufficient to balance the flow-induced torque. As cylinder velocity increases, the flow-induced torque also increases. Once this torque exceeds the maximum restoring torque generated by bottom-heaviness, a steady orientation can no longer be maintained, and the egg revolves around its center. In this regime, both the internal and external torques vary with egg orientation, producing periodic relative angular velocity. Indeed, the model predicted the observed frequency and amplitude well (Fig. 3D and Fig. 4A) with *R*^2^ = 0.95 for the frequency and RMSE of 7.3% of the mean measured amplitude. The estimated regression slopes did not differ significantly from 1, indicating low model bias.

### Younger eggs show greater stability across cylinder velocities

The size, position, and density of the oil globule, embryo, and remaining internal egg fluid change across developmental stages. Most dominant being the embryo elongation towards the upper hemisphere of the egg. These changes, in turn, alter the distance to the centers of mass, which has a central role in determining the magnitude of the egg’s gravitational torque, and thus its ability to stabilize itself. This effect if termed the reorientation time scale, B, in Durham et al. [2009]. While the kinematics of 1 DPF eggs were similar to those of 2 DPF eggs, our results suggest that the gravitational restoring torque (smaller B) of 1 DPF eggs was higher than that of 2 DPF eggs. It is evident in Fig. 3A-B where 1 DPF eggs maintain steady orientation at higher cylinder velocities compared to 2 DPF eggs. The mean angle of 1 DPF eggs is lower under the same flow conditions (see Fig. S1A). Also, their mean frequency was lower and mean amplitude higher than 2 DPF eggs (Fig. S1B-C). We used the model to predict egg behavior as a function of developmental stage and cylinder angular velocity (Fig. 4). The model predicted that the cylinder angular velocity at which the transition from the stable-orientation to revolving eggs occurred decreased with egg age. Below that transition speed, *ω*_rel_ depended only on cylinder velocity. Above this transition, *ω*_rel_ increased only with egg age.

## Discussion

Our results show that eggs of *S. aurata* do not behave as passive tracer particles in cylinderdriven flow. Bottom-heavy eggs exhibited rotational motion relative to the surrounding water under flow conditions corresponding to shear rates generated by turbulence in the ocean upper mixed layer and estuaries. This is important because turbulence consists of eddies, rotating structures of different scales, superimposed on the mean flow. While fish eggs are transported through the ocean by the mean flow, they also interact with eddies at the scale of the egg, which impose local rotational forcing. The magnitude of this relative motion depended on the imposed flow strength and developmental stage: early-stage eggs were more stable, whereas later-stage eggs transitioned to revolving motion at weaker flows. The egg’s dynamics were reproduced by a rigid-body model in which the egg’s bottom-heaviness shifts the center of mass relative to the egg’s center. This offset generates a gravitational restoring torque that competes with the external flow-induced torque. When the restoring torque is sufficient to balance the flow-induced torque, the egg maintains a stable orientation; and when the flow-induced torque exceeds the maximum restoring torque, the egg revolves around its center.

Turbulence shapes planktonic processes such as encounter rates, aggregation, and dispersal, but its effects on fish-egg development remain poorly understood. Rotation alone, however, does not necessarily increase oxygen exchange. At the small Reynolds numbers examined here, rotation moves water mainly along the egg surface. If oxygen consumption were evenly distributed, oxygen concentrations would vary only with distance from the egg, and this surface-parallel flow would not carry oxygen across that gradient. In real eggs, however, consumption is concentrated in the embryo, likely creating an uneven oxygen distribution. Studies of spheres with localized absorbing regions show that the size and position of these regions affect both the surrounding concentration pattern and the overall uptake rate **???**. Flow along the egg surface could therefore redistribute oxygen-rich water and potentially increase exchange. Testing this possibility will require models that combine water movement outside the egg, oxygen transport through the egg envelope and interior, localized consumption by the embryo, and changing egg–water rotation—a combination not addressed by existing studies.

Because oxygen availability constrains embryo development and survival, this unresolved coupling between flow and transport may have important ecological consequences. Yet small-scale flow conditions are rarely reported in respiration experiments or incorporated into ecological models. Measurements from mixed chambers may therefore include unrecognized flow-dependent transport, while models that ignore spatial and temporal variation in turbulence may misestimate oxygen supply and waste removal in the ocean.

Bottom heaviness has been considered an ecologically-relevant organisms Durham et al. [2009], DiBenedetto et al. [2025], Kessler [1985], Zeng et al. [2022], however it has been largely overlooked for non-motile ones. A dense embryo localized near the egg margin is present in the eggs of virtually all marine fish species. Additionally, at least one oil globule is present in 75% of marine fish eggs Ahlstrom and Moser [1980], suggesting that bottom-heaviness may be common and that relative motion may be a general attribute of many pelagic eggs. This interpretation is supported by observations in several teleost species showing that freely floating eggs often orient with the oil globule uppermost and the embryo below it Cambalik et al. [1998], Hildebrand and Cable [1930]. Bottom-heaviness may also arise in the absence of an oil globule, when a localized dense embryo is balanced by a low-density egg yolk or other internal structures. In fact, our study implies that any organism at these scales with a non-uniform distribution of mass may experience relative motion under rotational flows.

Beyond mass transfer, maintaining orientation relative to gravity may itself be important during embryogenesis. In many species, gravity provides spatial cues for early developmental patterning, including the establishment of body axes. Experiments on zebrafish and medaka eggs have shown that disrupting normal orientation can alter dorsoventral development or compromise bilateral symmetry Wacker et al. [1994], Fluck et al. [1998]. More recently, the formation of connecting tissue between the oil globule and embryo was shown to depend on egg orientation and to be important for larval survival Nande et al. [2024]. These observations suggest that both stability relative to gravity and relative motion that enhances exchange may be important, but their relative benefits may differ across developmental stages.

We therefore hypothesize that pelagic eggs may experience an optimal range of turbulence, in which enhanced mass transfer is balanced against the need to maintain steady orientation, particularly during early development. Our findings are relevant both to natural populations and to mariculture. Small-scale flow conditions may influence egg development in the ocean and could potentially be optimized in culture systems to enhance oxygen supply without disrupting steady orientation. Future measurements of embryo respiration and survival should therefore report and, where possible, control the flow conditions experienced by the eggs.

### Study system and experimental overview

We used Gilthead sea bream (*Sparus aurata*) as a model because they are representative marine pelagic fish whose eggs develop in the water column. Eggs were 0.995 *±* 0.0211 mm (n = 341) in diameter, spherical, and contained a visible oil globule with a diameter of 0.2014±0.001 mm (CITE Kamachi). Eggs were obtained approximately six hours after fertilization from a commercial hatchery in Eilat and maintained at controlled temperature of 16–17°C. According to the hatchery’s protocol, eggs were kept in filtered salinity-reduced seawater with salinity of 19 gr / liter. At this temperature, hatching is reported to occur between 51 and 53 hours post-fertilization Kamaci et al. [2005]. Accordingly, experiments were conducted over the two days of development (1 and 2 days post-fertilization, DPF), enabling comparison of egg behavior across developmental stage.

Experiments were conducted in a transparent plexiglass cylinder (internal diameter 32.1 mm and length 18.5 mm) mounted horizontally on a motorized rotation system, Fig 5. The cylinder was rotated at controlled angular velocities using a step motor, generating a reproducible rigid-body flow. Particle image velocimetry (PIV) showed a strong correspondence with rigid-body rotation across the tested speed range (see SI), and PIV-derived angular velocities closely matched the cylinder angular velocity. We recorded the egg’s motion and orientation using a high-speed video camera operating at 250 frames *s*^−1^ against a backlight illumination. Two frames of references were established for the analysis (see SI). In the cylinder frame of reference, the origin was set at the cylinder center, allowing egg position to be described as orbital motion relative to the cylinder. All analyzed eggs were advected with the solid body rotation of the water in the cylinder, maintaining an approximately constant radial position as they moved around the cylinder axis, Fig. 1a-d, f-i. Thus, the relative motion analyzed here refers to rotation of the egg around its own center. In the egg frame of reference, the egg center served as the origin, and the egg’s orientation angle, *ϕ*_egg_ (see Fig. 2) was defined as the angle between the vector connecting the oil globule to the egg’s geometric center and the positive *x*_2_ axis (the horizontal plane in the earth-fixed reference frame).

**Figure 5.**
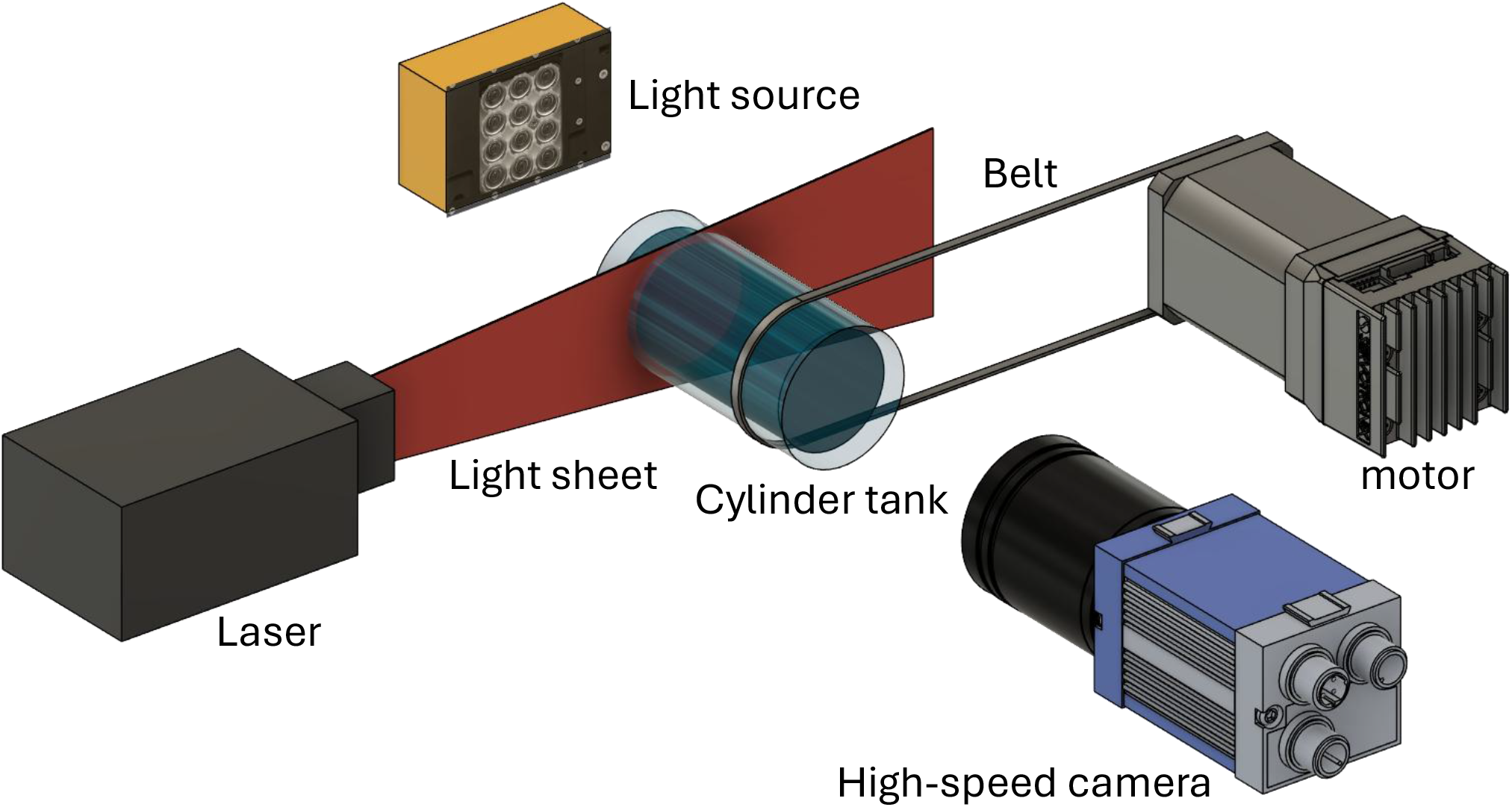
Schematic illustration of the experimental setup. A cylindrical tank, inner diameter 32.1 mm and length 18.5 mm, was filled with sea water and rotated at constant angular velocities using stepper motor. The motion of neutrally buoyant fish eggs inside the tank was recorded with a high-speed camera under backlight illumination. The flow field was measured using particle image velocimetry (PIV), with a laser sheet introduced perpendicular to the camera view.

Before each trial, seawater salinity in the cylinder was adjusted to obtain an approxi-mately neutral buoyancy of the eggs. Approximately 10 eggs were inserted into the cylinder, and tracked under thirteen cylinder angular velocities ranging 32.6–144.2 *deg s*^−1^. In total, we recorded 138 eggs from 74 videos (mean duration 38 *±* 7 *s* long) for 1 DPF eggs, and 115 videos for 2 DPF eggs. Videos were processed automatically, with 1,000 frames evenly spaces along the analyzed video. In each frame, egg outlines were detected in MATLAB using a Circular Hough Transform, which identifies circular objects and returns the radii and the coordinates of the egg’s center. The oil globule centers were detected using SLEAP Pereira et al. [2022], pre-trained on manually labeled images. The two outputs were merged by assigning each detected oil globule to an egg when the globule center fell within the corresponding egg boundary, accounting for the oil-globule radius. Cases with ambiguity, such as overlapping eggs or conflicting assignments, were excluded, as were tracks shorter than 500 frames. Detections with implausible egg diameters were removed using a size-based threshold around the observed egg-diameter distribution. Tracks in which the egg center did not maintain a constant radial distance from the cylinder center were also excluded. Finally, eggs that exhibited rotation around out-of-plane axes in addition to rotation around the main cylinder axis were excluded from further analysis. These cases represented a minority of the observations (26%); because the measurements were two-dimensional, such multi-axis rotation could not be accurately resolved or quantified.

### Mathematical model of the egg rotation

To interpret the experiments, we developed an analytical rigid-body model for egg rotation using Euler’s equation, see Box. 1. To obtain the values for the center of mass position vector, **d**, and moment of inertia **I** the egg’s internal structure was represented as a three-component composite body: embryo, oil globule, and internal fluid (yolk + perivitelline fluid). The embryo was approximated as a partial torus, the oil globule as a sphere, and the remaining volume as fluid. These geometries were parameterized based on images of eggs from the two ages obtained from our experiments. Centers of mass and moments of inertia were computed for each component and then combined to obtain the whole-egg center of mass and inertia tensor. Some parameters were measured or taken from literature, while others were treated as variable inputs (e.g., embryo geometry over development, external forcing) For the external torque, we assumed low Reynolds number flow at the egg scale and used a superposition approach: rigid-body cylinder flow plus the flow around a rotating sphere.

The model also estimated the local shear scale showing that the experimental shear range overlaps the range expected in upper mixed-layer and estuarine habitats.

## Data Availability

To allow others to replicate and build on work published in PNAS, authors must make materials, data, and associated protocols, including code and scripts, available to readers in a public repository upon publication.

## Author Contributions

Conceptualization: MVG, US, RH. Methodology: all authors. Formal analysis: MVG. Investigation: MVG, US, RH. Writing – original draft preparation: MVG, RH; reviewing and editing – US, DG, DR. Supervision: US, RH.

## Acknowledgments

Please include your acknowledgments here.

## Box 1.

**Mathematical model of fish egg rotation.** The pelagic fish egg was modeled as a spherical rigid body. Its rotational dynamics were described using Euler’s equation in an egg-centered coordinate system whose axes rotate with the egg (Fig. 2):

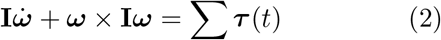

where **I** is the egg mass moment of inertia tensor,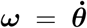 is the vector of the egg angular velocity relative to the earthbound frame, ***θ*** is the vector of the rotation angle and ***τ*** (*t*) is the vector of the torques acting on the egg. Two torques are acting on the egg under these conditions; external flow-induced torque, ***τ*** _*ext*_, and gravitational restoring torque, ***τ*** _*int*_.

To simplify the model, we restricted it to the exper-imentally resolved rotation about the *x*_1_ axis, and did not treat cases in which perturbations generate rotation about the other axes. The flow-induced torque was estimated using the expression, given by Lamb (1945), for the viscous torque on a sphere rotating in an otherwise quiescent fluid:

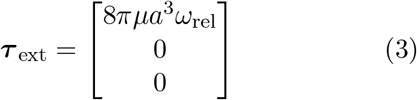

where *µ* is the water dynamic viscosity, *a* is the egg radius, and *ω*_rel_ is the water angular velocity relative to the egg (Eq. 1). The restoring torque arises from the offset, **d**, between the egg’s center (the origin in Fig. 2) and the center of mass (marked in yellow in Fig. 2),

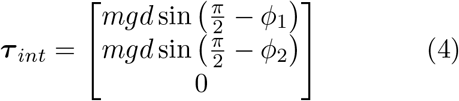

where *m* is the egg mass, *g* is the gravitational acceleration, and *ϕ*_1_ and *ϕ*_2_ are the oil globule orientation angles, with respect to *x*_1_ and *x*_2_ axes. Note that *ϕ*_1_ is *ϕ*_*egg*_ in Fig. 2. Using Eqs. 3 and 4, Eq. 2 is reduced to

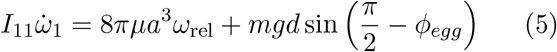

where 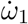 is the time derivative of −*ω*_rel_. The off-set between the egg center and center of mass, **d**, and the moment of inertia, **I**, were calculated from the size, position, and density of the oil globule, embryo, and remaining egg internal fluid. The detailed calculation is described in the SI.

