## Supplementary Information for "Bottom-heaviness stabilizes pelagic fish eggs and generates advective transport at their surface"

### Supporting Information: Non-uniform internal structure generates relative water motion around pelagic fish eggs in rotating flow

Maya Fanny van-Gelder, Uri Shavit, Dan Givoli, Daniel Rabinovich and Roi Holzman

#### 0.1 Methods

### Experimental setup and flow validation

#### Rotating-cylinder setup

Experiments were conducted in a transparent rotating cylinder designed to impose a controlled rotational flow on freely suspended fish eggs. The cylinder had an internal diameter of 32.1 mm and an internal chamber length of 18.5 mm. It was mounted horizontally and rotated by an electric step motor through a pulley-and-belt transmission. The imposed cylinder angular velocity is denoted by $\omega_{\text{cyl}}$.

Egg motion and water motion were recorded using two high-speed cameras (CP70-2-M/C-1000, Optronis GmbH, Germany; 1920 × 1080 pixels), each equipped with a 105 mm NIKKOR lens and positioned at opposite ends of the cylinder. One camera recorded the eggs under backlight illumination, while the second camera recorded tracer particles for particle image velocimetry (PIV). Images taken by the egg camera were illuminated by a synchronized strobe light, whereas images taken by the PIV camera were illuminated by a continuous-wave laser sheet passing through the center of the cylinder. Hollow glass spheres of 10 $\mu$m diameter were added as PIV tracer particles. For both cameras each pixel was equivalent to 0.025 mm.

Before each experiment, eggs were inserted into the cylinder and the salinity was adjusted to obtain neutral buoyancy. The cylinder was then rotated at prescribed angular velocities, and the orientation and motion of the eggs were recorded.

#### Coordinate systems and egg motion

Three Cartesian coordinate systems were used to describe the egg position, orientation, and internal geometry (Fig. S1). The cylinder-centered coordinate system, $(X_1, X_2, X_3)$, has its origin at the center of the cylinder in the imaging plane, with the $X_1$ axis aligned with the cylinder axis and the $X_2$-$X_3$ plane corresponding to the observed cylinder cross-section. The radial distance of the egg center from the cylinder center is denoted $R_{\mathrm{orb}}$.

A second coordinate system, $(x_1, x_2, x_3)$, was centered on the egg but remained aligned with the cylinder frame, such that $x_1 \parallel X_1$, $x_2 \parallel X_2$, and $x_3 \parallel X_3$. Its origin therefore translated with the egg as the egg center moved around the cylinder, while its axes remained fixed in orientation relative to the laboratory frame.

A third, body-fixed coordinate system, $(x_1', x_2', x_3')$, was attached to the egg and rotated with it. The $x_1'$ axis coincides with $x_1$ and is normal to the imaging plane. The positive $x_3'$ axis was defined from the egg center toward the oil-globule center, and $x_2'$ was defined perpendicular to $x_3'$ in the imaging plane. The egg orientation angle, $\phi_{\mathrm{egg}}$, was defined as the angle between the positive $x_2$ and $x_3'$ directions.

The body-fixed frame was used to describe the internal egg geometry and calculate the mass properties. The geometric parameters used in the rotational-dynamics model, including the egg radius $a$, oil-globule radius $a_{\mathrm{oil}}$, embryo major radius $r_{\mathrm{emb}}$, embryo minor radius $a_{\mathrm{emb}}$, and embryo angular extent $\alpha$, are defined in Fig. S1.

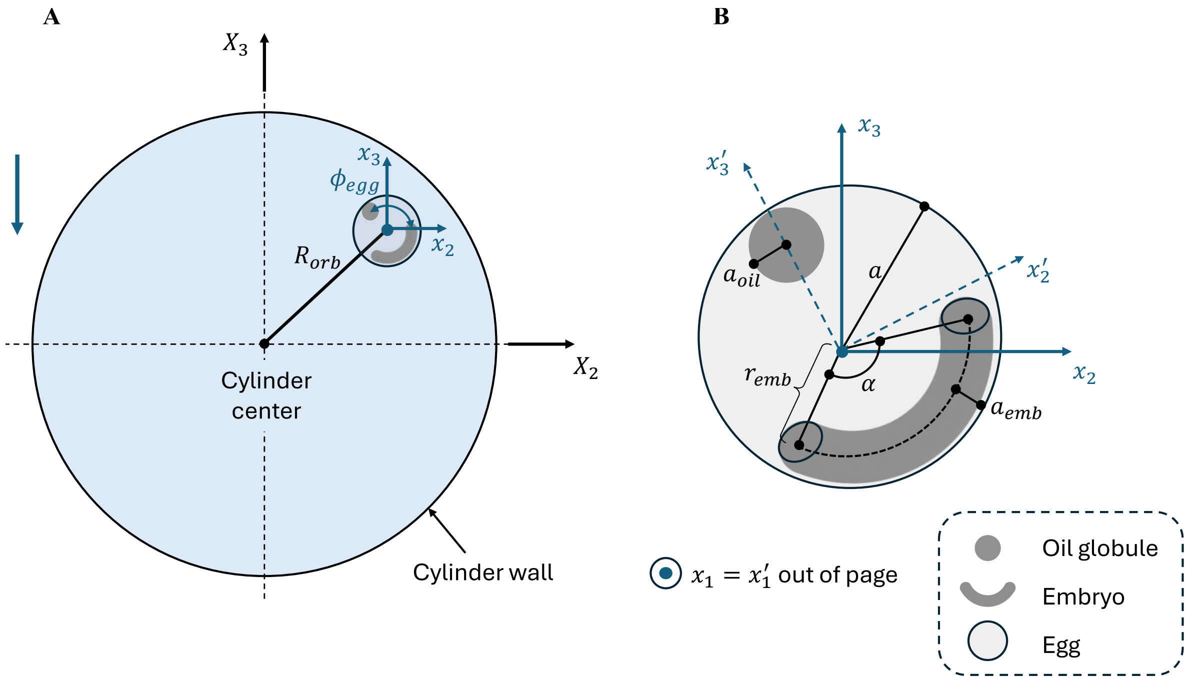


Figure S1: Coordinate systems and geometric parameters used in the analysis. (A) Cylinder-centered coordinate system viewed along the $X_1$ axis and the local egg-centered coordinates. $R_{orb}$ is the egg distance from the center of the rotating cylinder, and $\phi_{egg}$ is the egg orientation. (B) Egg-centered coordinate system and geometric parameters used in the rotational-dynamics model, including the egg radius $a$, oil-globule radius $a_{\text{oil}}$, embryo major radius $r_{\text{emb}}$, embryo minor radius $a_{\text{emb}}$, and embryo angular extent $\alpha$. The illustration is schematic and not drawn to scale.

### PIV validation of solid-body rotation

Particle image velocimetry (PIV) analysis was performed using a modified version of the open-source MATLAB toolbox MatPIV v. 1.6.1 Sveen [2004]. For each cylinder angular velocity, 1,000 image pairs from one video were analyzed using 32 × 32 pixel interrogation windows with 50% overlap, resulting in a velocity-vector spacing of 0.397 mm. The time separation between paired images was selected separately for each imposed angular velocity to obtain a particle displacement of approximately six pixels; because the images were recorded at 250 fps, the minimum available time separation was 4 ms. Velocity vectors were filtered using a local median criterion with a threshold of 2.8 times the standard deviation of the eight neighboring vectors. Approximately 0.07% of the vectors were rejected by this criterion and replaced by interpolation. Regions outside the internal cylinder chamber were masked before the PIV analysis.

PIV was used to validate the flow field inside the rotating cylinder. The analysis had two purposes: first, to test whether the water motion in the absence of eggs could be approximated as solid-body rotation; and second, to verify that the cylinder angular velocity calculated from the motor input could be used as $\omega_{\mathrm{cyl}}$.

The time-averaged velocity magnitude, $U_{\mathrm{PIV}}$, at each PIV location was calculated as

$$U_{\mathrm{PIV}} = \sqrt{U_{2,\mathrm{PIV}}^2 + U_{3,\mathrm{PIV}}^2} \tag{S1}$$

where $U_{2,\mathrm{PIV}}$ and $U_{3,\mathrm{PIV}}$ are the measured velocity components in the PIV image plane. For solid-body rotation, the velocity magnitude is expected to increase linearly with radial distance from the rotation center according to

$$U_{\mathrm{PIV}} = \omega_{\mathrm{PIV}} \sqrt{(X_{2,\mathrm{PIV}} - X_{2,0})^2 + (X_{3,\mathrm{PIV}} - X_{3,0})^2}, \tag{S2}$$

where $X_{2,\mathrm{PIV}}$ and $X_{3,\mathrm{PIV}}$ are the corresponding image-plane coordinates. The parameters $X_{2,0}$, $X_{3,0}$, and $\omega_{\mathrm{PIV}}$ were estimated simultaneously by fitting Eq. S2 to the measured time-averaged velocity field. Thus, $X_{2,0}$ and $X_{3,0}$ define the estimated center of rotation, while $\omega_{\mathrm{PIV}}$ is the corresponding angular velocity.

Across all tested cylinder angular velocities, the mean coefficient of determination for the relationship between $U_{\mathrm{PIV}}$ and radial distance was $R^2 = 0.904$, supporting the approximation of the measured flow as solid-body rotation. The angular velocities obtained independently from the PIV measurements, $\omega_{\mathrm{PIV}}$, closely followed the corresponding cylinder angular velocities calculated from the motor input (Fig. S@@). Therefore, in all subsequent analyses, $\omega_{\mathrm{cyl}}$ denotes the cylinder angular velocity calculated from the motor input.

### Environmental shear-scale comparison

The cylinder angular velocities used in the experiments were chosen to represent the range of shear rates experienced by pelagic fish eggs in their natural environment. In the upper mixed layer and in estuaries, where most pelagic fish eggs are found, turbulent eddies at the scale of the egg impose local rotational forcing. The turbulent shear rate in these environments typically ranges from approximately $4 \times 10^{-3}$ to 10 $s^{-1}$ Dillon and Caldwell [1980], Song and Rau [2022].

To compare the experimental flow conditions with this environmental range, we estimated the shear scale at the egg surface. As an upper bound, we assume the egg remains stationary while the surrounding fluid rotates at $\omega_{\mathrm{cyl}}$, giving a maximum tangential velocity of the fluid relative to the egg surface of

$$u_{\phi,\max} = \omega_{\mathrm{cyl}} a, \tag{S3}$$

where $a = 0.5$ mm is the egg radius. The corresponding shear scale at the egg surface is estimated as

$$\dot{\gamma}_{\max} \sim \frac{u_{\phi,\max}}{a} = \omega_{\mathrm{cyl}}. \tag{S4}$$

For the cylinder angular velocities used in the experiments (32.56–144.19° $s^{-1}$, corresponding to 0.57–2.52 rad $s^{-1}$), this gives a shear-scale range of 0.57–2.52 $s^{-1}$. This range overlaps with the upper portion of the environmental shear range, corresponding to moderately to highly energetic conditions in the upper mixed layer and estuaries. The validity of the upper-bound assumption in Eq. S3 is discussed in SI section @@ (Flow-induced torque), where the flow field around the egg is derived explicitly.

### Egg tracking and kinematic analysis

#### Egg and oil-globule detection

To obtain the egg orientation over time, the egg surface and oil globule were identified in each analyzed video frame. The egg surface provided the egg center and radius, while the oil globule position relative to the egg center was used to calculate the egg orientation angle $\phi_{egg}$ (see SI section @@ for coordinate system definitions).

The oil globule center was detected using SLEAP Pereira et al. [2022], an open-source deep-learning framework for multi-animal pose tracking. A single node was defined per egg, located at the center of the oil globule. The model was trained on 2,804 manually labeled egg instances across 301 frames, drawn from one 1 DPF and one 2 DPF video. Of these, 90% (271 frames) were used for training and 10% (30 frames) for validation, using the top-down multi-animal method. Once trained, SLEAP detected and linked oil-globule positions across successive frames. The output was the estimated center coordinates of each oil globule in each analyzed frame.

The egg surface was detected using a MATLAB implementation of the Circular Hough Transform (CHT) Davies [2005], which identifies circular objects from image edges. The output consisted of the estimated center coordinates and radius of each detected egg.

The mean video duration was 38.6 s, corresponding to 9,640 frames recorded at 250 fps. To reduce computational cost, 1,000 frames were evenly sampled from each video, giving an effective frame rate of approximately $25 \pm 1.8$ fps. This sampling rate was sufficient for subsequent tracking and kinematic analysis.

The SLEAP and CHT outputs were combined by assigning an oil globule to an egg when the distance between their detected centers was less than or equal to the egg radius minus the oil globule radius. To avoid ambiguous assignments, oil globules that could be associated with more than one egg, and eggs with more than one candidate oil globule, were excluded.

#### Calculation of egg orientation and relative angular velocity

The orientation angle $\phi_{\text{egg}}$ was defined as the angle between the vector from the egg center to the oil globule center and the positive $x_2$ axis, expressed over 0 deg $\leq \phi_{\text{egg}} <$ 360 deg.

The orientation time series was unwrapped to remove discontinuities at 360 deg, producing a continuous angle record $\widetilde{\phi}_{\text{egg}}(t)$. The egg angular velocity was then estimated from the unwrapped angle using a centered finite difference with a 30-frame separation:

$$\omega(t_i) = \frac{\widetilde{\phi}_{\text{egg}}(t_{i+15}) - \widetilde{\phi}_{\text{egg}}(t_{i-15})}{t_{i+15} - t_{i-15}}. \tag{S5}$$

The relative angular velocity was then obtained as

$$\omega_{\text{rel}}(t_i) = \omega_{\text{cyl}} - \omega(t_i), \tag{S6}$$

where $\omega_{\text{cyl}}$ is the cylinder angular velocity, as defined and validated by PIV above.

#### Track filtering and validation

The automated detection was followed by a two-stage filtering procedure: first at the detection level, to remove likely misidentifications, and then at the track level, to remove unreliable trajectories. A final validation step was used to confirm that the accepted tracks were physically consistent with the imposed flow.

**Diameter-based filtering.** The natural variability in *S. aurata* egg diameter is lower than 3% Rønnestad et al. [1994], Kamaci et al. [2005]. Detections with diameters deviating by more than two standard deviations from the measured mean egg diameter (0.988±0.034 mm, corresponding to ±6.8%) were flagged as likely misidentifications and excluded. Manual inspection of a sample of excluded detections confirmed that the causes included misidentification of dirt or hatched larvae as eggs, incorrect detection during early developmental stages when the gastrulation cap disrupted circular edge detection by the CHT algorithm, and overlapping eggs (Fig. S@@). Following this filter, 4.19% of detections were removed, and the mean egg diameter of the accepted detections was 0.990 mm.

**Track-based filtering.** Tracks containing 500 detections or fewer were excluded as predominantly representing low-quality detections. For longer tracks, a radial-consistency criterion was applied to ensure that each track represented a single egg rather than jumping between eggs. Since eggs move in approximately circular orbits around the cylinder axis, the radial distance from the cylinder center, $R_{\mathrm{orb}}$, should remain approximately constant within a track. Tracks in which $R_{\mathrm{orb}}$ deviated by more than 0.692 mm (25 pixels) from the track mean were rejected (Fig. S@@).

**Validation.** The filtering methodology was validated by comparing the imposed cylinder angular velocity, $\omega_{\mathrm{cyl}}$, with the orbital angular velocity of the egg center,

$$\omega_{\mathrm{orb}} = \frac{360\,deg}{T_{\mathrm{orb}}}, \tag{S7}$$

where $T_{\mathrm{orb}}$ is the mean time required for the egg center to complete one orbit around the cylinder axis. The close agreement between $\omega_{\mathrm{orb}}$ and $\omega_{\mathrm{cyl}}$ across the tested velocities (Fig. S@@) confirmed that the accepted eggs were advected with the solid-body rotation of the water.

#### Classification of steady-oriented and revolving eggs

Accepted tracks were first classified according to the total angular range spanned by $\phi_{\mathrm{egg}}$ over the track duration. The angular range was defined as

$$\Delta\phi_{\mathrm{egg}} = \phi_{\mathrm{egg,max}} - \phi_{\mathrm{egg,min}}. \tag{S8}$$

Tracks with $\Delta\phi_{\mathrm{egg}} < 50\,deg$ were classified as steady-oriented eggs, in which the egg maintains a near-constant orientation (Vid. S@@). Tracks with $\Delta\phi_{\mathrm{egg}} > 300\,deg$ were classified as revolving eggs, in which the egg completed at least one full rotation about the resolved $x_1$ axis (Vid. S@@).

**Exclusion of multi-axis rotation.** Among the accepted tracks, those with an angular range (maximum minus minimum $\phi_{\mathrm{egg}}$) between $50\,deg$ and $300\,deg$ could not be unambiguously classified as either steady-oriented or revolving. Manual inspection of a random sample of 35 such tracks showed apparent out-of-plane motion consistent with rotation about axes in addition to $x_1$. Since the measurements were performed in 2D, such multi-axis rotation could not be accurately resolved and quantified. All tracks within this intermediate group were therefore excluded from further analysis. In total, 150 eggs were excluded on this basis.

However, automatic classification based on angular range alone was insufficient to exclude all multi-axis rotation cases. Manual inspection of all tracks classified as revolving ($\phi_{\mathrm{egg,max}} - \phi_{\mathrm{egg,min}} > 300\,deg$) revealed that some tracks within this group also exhibited multi-axis rotation. An additional 90 tracks were excluded following this manual review, leaving a final set of revolving egg tracks for further analysis.

### Calculation of revolution frequency and relative angular velocity amplitude

For each track classified as revolving, the unwrapped egg-orientation record, $\widetilde{\phi}_{\mathrm{egg}}(t)$, was fitted over the complete track using

$$\widetilde{\phi}_{\mathrm{egg}}(t) = \phi_0 + \Omega_{\mathrm{egg}} t, \tag{S9}$$

where $\Omega_{\mathrm{egg}}$ is the mean fitted angular velocity of the egg over the track. The egg revolution frequency was then calculated as

$$f_{\mathrm{egg}} = \frac{\Omega_{\mathrm{egg}}}{360\,deg}, \tag{S10}$$

with $f_{\mathrm{egg}}$ expressed in Hz when $\Omega_{\mathrm{egg}}$ is expressed in degrees per second.

To calculate the relative-angular-velocity amplitude, the dominant cycle duration was estimated from the autocorrelation of the smoothed $\omega_{\mathrm{rel}}(t)$ signal and used to identify successive cycles. Within each cycle, the signal was linearly detrended and median-filtered, and a robust peak-to-peak range was calculated as the difference between its 95th and 5th percentiles. The reported relative-angular-velocity amplitude, $A_\omega$, was defined as half the mean robust peak-to-peak range across all detected cycles.

#### Error estimation

To assess the accuracy of the automatically calculated orientation angle, $\phi_{\text{egg}}$, manual measurements were performed on a subset of 62 accepted egg tracks: 30 tracks of 1 DPF eggs and 32 tracks of 2 DPF eggs. For each track, five points were manually selected in one representative frame: one at the center of the oil globule and four along the egg capsule. A circle was fitted to the four peripheral points to determine the egg center and radius, after which $\phi_{\text{egg}}$ was calculated using the same geometric definition as for the automated measurements.

The manually measured angles were compared with the corresponding automated values. For 1 DPF eggs, the mean absolute error was $4.10 \pm 2.63\,deg$, and the mean signed error, defined as automated minus manual angle, was $+1.05\,deg$. For 2 DPF eggs, the mean absolute error was $5.67 \pm 5.24\,deg$, and the mean signed error was $+0.48\,deg$. The small positive signed errors indicate only a weak systematic tendency for the automated method to overestimate $\phi_{\text{egg}}$.

#### Embryo morphology measurements

In addition to the kinematic quantities described above, embryo morphology was manually measured for use in the rotational-dynamics model. For each egg, up to 40 candidate frames were sampled at approximately uniform intervals over the tracked sequence and inspected until eight suitable frames had been measured. In each accepted frame, two points were placed at the visible ends of the embryo, and two additional points were placed across its transverse thickness. The measured point coordinates were expressed relative to the egg center in the egg-centered imaging frame $(x_1, x_2, x_3)$. Assuming that the embryo ends lay on the spherical egg surface of radius $a$, the out-of-plane coordinate of each endpoint was reconstructed as

$$x_1 = \pm\sqrt{\max(0, a^2 - x_2^2 - x_3^2)}.$$

The two end points were assigned to opposite hemispheres, and their three-dimensional angular separation was calculated as

$$\tilde{\alpha} = \cos^{-1}\left(\frac{\mathbf{r}_1 \cdot \mathbf{r}_2}{|\mathbf{r}_1||\mathbf{r}_2|}\right) \tag{S11}$$

where $\mathbf{r}_1$ and $\mathbf{r}_2$ are the reconstructed position vectors of the two embryo endpoints relative to the egg center. Because the visible endpoint separation corresponded to different embryo-covered arcs at the two developmental stages, the embryo angular extent was defined as

$$\alpha = \begin{cases} \tilde{\alpha}, & 1 \text{ DPF}, \\ 2\pi - \tilde{\alpha}, & 2 \text{ DPF}. \end{cases} \tag{S12}$$

The transverse embryo thickness in each frame was calculated as the Euclidean distance between the two thickness points, and the embryo minor radius, $a_{\text{emb}}$, used in the model was half this distance. The reported values of $\alpha$ and $a_{\text{emb}}$ were obtained from the mean over eight accepted frames, and the corresponding between-frame standard deviations were retained as estimates of measurement variability.

Early developmental stages, before formation of a distinct embryo, were excluded from figure panels and model comparisons requiring the partial-torus embryo geometry. Eggs for which eight suitable frames could not be obtained from the candidate pool were classified as unmeasurable and excluded from analyses requiring embryo morphology. They were retained in analyses of the proportions of observed behavioral classes, which depended only on track duration and observed angular range.

### Model development

#### Model overview and assumptions

The egg was modeled as a neutrally buoyant rigid sphere with a nonuniform internal mass distribution. Its rotational dynamics were described by Euler's equation for rigid body rotation. The model accounts for the balance between two torques: a flow-induced torque arising from the relative angular velocity between the surrounding water and the egg, and a gravitational restoring torque arising from the offset between the egg center and the egg center of mass. Viscous resistance was represented entirely by the flow induced torque that will be discussed in the next chapter. The model was restricted to rotation about the experimentally resolved $x_1$ axis. Values of all parameters used in the model are summarized in Table S1.

#### Governing equation and one-axis reduction

The pelagic fish egg was modeled as a rigid body with a non-uniform internal mass distribution. Its rotational dynamics were described by Euler's equation, expressed in the body-fixed coordinate system $(x_1', x_2', x_3')$ defined in Fig. S1:

$$\mathbf{I}\dot{\boldsymbol{\omega}} + \boldsymbol{\omega} \times (\mathbf{I}\boldsymbol{\omega}) = \boldsymbol{\tau}_{\mathrm{ext}}(t) + \boldsymbol{\tau}_{\mathrm{int}}(t), \tag{S13}$$

where $\mathbf{I}$ is the egg mass moment-of-inertia tensor, $\boldsymbol{\omega}$ is the angular velocity of the egg relative to the laboratory frame expressed in the body-fixed coordinates, and $\boldsymbol{\tau}_{\mathrm{ext}}$ and $\boldsymbol{\tau}_{\mathrm{int}}$ are the flow-induced and gravitational torques, respectively, expressed in the same frame. The body-fixed axes were chosen along the principal axes of inertia, such that $\mathbf{I}$ is constant and diagonal in this coordinate system.

The model was restricted to the experimentally resolved rotation about the $x_1'$ axis, which coincides with $x_1$ and is normal to the imaging plane. Rotation about the other two axes was not considered. Under this one-axis restriction, the rotation angle about $x_1'$, $\phi_1$, was identified with the measured egg orientation angle,

$$\phi_1 \equiv \phi_{\text{egg}}, \qquad \omega = \dot{\phi}_1 = \dot{\phi}_{\text{egg}}. \tag{S14}$$

Because the rotation is restricted to a principal axis, the gyroscopic cross-product term vanishes, and Eq. S13 reduces to

$$I_{11}\dot{\omega} = \tau_{\text{ext},1}(t) + \tau_{\text{int},1}(t), \tag{S15}$$

where $I_{11}$ is the egg moment of inertia about the $x_1'$ axis.

The flow-induced torque, $\tau_{\text{ext},1}$, gravitational restoring torque, $\tau_{\text{int},1}$, and moment of inertia, $I_{11}$, are defined below.

### Flow-induced torque

The flow-induced torque was approximated using the Stokes-flow solution for a small sphere rotating in an otherwise quiescent fluid. In the absence of the egg, the rotating-cylinder flow was solid-body rotation (as validated by PIV measurements, see SI section *PIV validation of solid-body rotation*). In the egg-centered laboratory-aligned frame, the surrounding solid-body flow rotates about $x_1$ with angular velocity $\omega_{\text{cyl}}$, while the egg rotates about the same axis with angular velocity $\omega$. The angular velocity of the surrounding water relative to the egg is therefore

$$\omega_{\text{rel}} = \omega_{\text{cyl}} - \omega. \tag{S16}$$

Because $x_1' = x_1$, this relative angular velocity has the same scalar component along the $x_1'$ axis of the body-fixed coordinate system used in Euler's equation.

For a sphere at low Reynolds number the flow-induced torque about $x_1'$ is **?**,

$$\tau_{\mathrm{ext},1} = 8\pi\mu a^3 \omega_{\mathrm{rel}}, \tag{S17}$$

where $a$ is the egg radius and $\mu$ is the dynamic viscosity.

The egg-scale Reynolds number was estimated using the maximum tangential velocity at the egg surface. The cylinder angular velocities used in the experiments were 32.56–144.19° $\mathrm{s}^{-1}$, corresponding to

$$u_{\phi,\max} = \omega_{\mathrm{cyl}} \frac{\pi}{180} a = 0.284\text{–}1.26 \ \mathrm{mm\ s^{-1}}. \tag{S18}$$

Using the egg diameter, $2a$, as the characteristic length scale, the corresponding Reynolds number was

$$Re = \frac{\rho_w u_{\phi,\max}(2a)}{\mu} = 0.268\text{–}1.19, \tag{S19}$$

where $\rho_w$ is the seawater density. The resulting range, $Re = 0.268\text{–}1.19$, is within the low-Reynolds-number regime in which viscous stresses dominate the rotational forcing.

For revolving eggs, $\omega_{\mathrm{rel}}$ varies over time, whereas the classical Stokes–Lamb solution is derived for steady rotation of a sphere. We therefore used this solution as a quasi-steady approximation, in which the instantaneous flow-induced torque is assumed to be determined by the instantaneous relative angular velocity between the water and the egg. The validity of this approximation was characterized by comparing the timescale of temporal changes in the azimuthal flow with the timescale of viscous momentum diffusion. Using $u_\phi \sim a|\omega_{\mathrm{rel}}|$, their ratio scales as

$$\frac{\partial u_\phi/\partial t}{\nu \nabla^2 u_\phi} \sim \frac{a^2|\dot{\omega}_{\mathrm{rel}}|}{\nu|\omega_{\mathrm{rel}}|}, \tag{S20}$$

where $\nu = \mu/\rho_w$ is the kinematic viscosity of seawater. In the case producing the largest observed angular acceleration, this ratio was approximately 0.28. Thus, even under the most rapidly varying measured conditions, the unsteady contribution remained substantially smaller than the viscous contribution, supporting the quasi-steady approximation.

#### Gravitational restoring torque

The gravitational restoring torque arises from the offset between the egg geometric center and its center of mass. The vector from the egg center to the center of mass is denoted by $\mathbf{d}$ and is expressed in the body-fixed coordinate system. Because the internal mass distribution is fixed relative to the egg, $\mathbf{d}$ is constant in this frame.

The gravitational acceleration vector, $\mathbf{g}$, has a fixed direction in the laboratory frame, while its components in the body-fixed frame vary with the egg orientation. The egg mass was calculated from neutral buoyancy as

$$m = \frac{4}{3}\pi a^3 \rho_w. \tag{S21}$$

The gravitational torque about the egg center is

$$\boldsymbol{\tau}_{\text{int}} = \mathbf{d} \times m\mathbf{g}. \tag{S22}$$

For the modeled geometry, the center of mass lies along the $x_3'$ axis, so $d = |\mathbf{d}|$ denotes the distance between the egg center and its center of mass. For rotation about $x_1'$, the gravitational torque reduces to

$$\tau_{\text{int},1} = mgd \sin\left(\frac{\pi}{2} - \phi_{\text{egg}}\right). \tag{S23}$$

#### Geometric representation of the egg

The egg was represented as a neutrally buoyant rigid sphere of radius $a$ (Fig. S1). Within this sphere, the internal mass distribution was represented using three components: (i) a spherical oil globule of radius $a_{\text{oil}}$, (ii) an embryo represented as a partial torus, and (iii) the remaining volume, treated as internal fluid comprising the yolk and perivitelline fluid.

The internal geometry was defined in the body-fixed coordinate system $(x_1', x_2', x_3')$. The positive $x_3'$ axis was directed from the egg center toward the oil-globule center, with the embryo located on the opposite side of the egg. Because this coordinate system rotates with the egg, the positions of the internal components remain fixed in this frame.

The oil globule was assumed to be tangent to the inner surface of the egg. Its center was therefore located at

$$\mathbf{r}'_{\text{oil}} = \begin{bmatrix} 0 \\ 0 \\ a - a_{\text{oil}} \end{bmatrix}, \tag{S24}$$

relative to the egg center.

The embryo was represented as a partial torus in the $x_2'$–$x_3'$ plane. The torus was described by a minor radius $a_{\text{emb}}$, a major radius

$$r_{\text{emb}} = a - a_{\text{emb}}, \tag{S25}$$

and an angular extent $\alpha$. The partial torus was positioned symmetrically about the negative $x_3'$ direction, opposite the oil globule. By symmetry, the $x_1'$ and $x_2'$ coordinates of the embryo centroid are zero, and its centroid lies along the $x_3'$ axis.

The corresponding component volumes are

$$V_{\text{egg}} = \frac{4}{3}\pi a^3, \tag{S26}$$

$$V_{\text{oil}} = \frac{4}{3}\pi a_{\text{oil}}^3, \tag{S27}$$

$$V_{\text{emb}} = 2\pi^2 r_{\text{emb}} a_{\text{emb}}^2 \frac{|\alpha|}{2\pi}. \tag{S28}$$

Thus, $V_{\text{emb}}$ is the volume of a complete torus multiplied by the fraction of the toroidal circumference occupied by the embryo, $\alpha/2a$.

#### Component densities and neutral buoyancy

Let $\rho_{\text{oil}}$, $\rho_{\text{emb}}$, and $\rho_f$ denote the densities of the oil globule, embryo, and remaining internal fluid, respectively.

The embryo density $\rho_{\text{emb}}$ was treated as a model parameter. The density $\rho_f$ of the remaining internal fluid was calculated by imposing neutral buoyancy. Neutral buoyancy requires the total egg mass to equal the mass of the displaced seawater:

$$\rho_f \left(V_{\text{egg}} - V_{\text{oil}} - V_{\text{emb}}\right) + \rho_{\text{oil}} V_{\text{oil}} + \rho_{\text{emb}} V_{\text{emb}} = \rho_w V_{\text{egg}}. \tag{S29}$$

Solving Eq. S29 for $\rho_f$ gives

$$\rho_f = \frac{\rho_w V_{\text{egg}} - \rho_{\text{oil}} V_{\text{oil}} - \rho_{\text{emb}} V_{\text{emb}}}{V_{\text{egg}} - V_{\text{oil}} - V_{\text{emb}}}. \tag{S30}$$

For the calculation of the center of mass and moment of inertia, the composite egg can equivalently be represented as a sphere of density $\rho_f$, together with density-contrast contributions from the oil globule and embryo:

$$\Delta\rho_{\text{oil}} = \rho_{\text{oil}} - \rho_f, \tag{S31}$$

$$\Delta\rho_{\text{emb}} = \rho_{\text{emb}} - \rho_f. \tag{S32}$$

The corresponding effective mass contributions are

$$\Delta m_{\text{oil}} = V_{\text{oil}} \Delta\rho_{\text{oil}}, \tag{S33}$$

$$\Delta m_{\text{emb}} = V_{\text{emb}} \Delta\rho_{\text{emb}}. \tag{S34}$$

These effective mass contributions may be negative, as for the oil globule because $\rho_{\text{oil}} < \rho_f$. They therefore represent algebraic density-contrast contributions used in the center of mass and moment of inertia calculations, rather than the physical masses of the components.

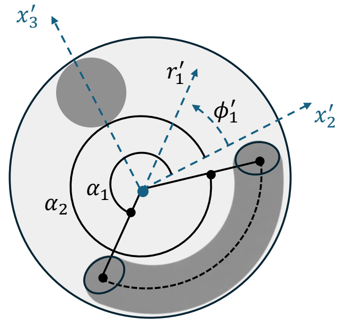


Figure S2: Geometric definition of the angular coordinate used to calculate the centroid of the partial-torus embryo in the body-fixed coordinate system.

#### Center of mass

The center-of-mass calculation was performed in the body-fixed coordinate system $(x_1', x_2', x_3')$. The embryo was modeled as a partial torus positioned symmetrically about the negative $x_3'$ axis. By symmetry, its centroid lies on the $x_3'$ axis, such that

$$r'_{\text{emb},1} = r'_{\text{emb},2} = 0.$$

The remaining centroid coordinate is

$$r'_{\text{emb},3} = \frac{1}{V_{\text{emb}}} \int_{V_{\text{emb}}} x_3' \, dV. \tag{S35}$$

Cylindrical coordinates $r_1'$ and $\phi_1'$ about the $x_1'$ axis were defined by (Fig. S2)

$$x_2' = r_1' \cos \phi_1', \qquad x_3' = r_1' \sin \phi_1'. \tag{S36}$$

The limits of the partial torus in the $\phi_1'$ direction were defined as $\alpha_1$ and $\alpha_2$ (Fig S2), such that

$$\alpha_2 - \alpha_1 = \alpha. \tag{S37}$$

By from symmetry

$$\alpha_1 = \frac{3\pi}{2} - \frac{\alpha}{2}, \qquad \alpha_2 = \frac{3\pi}{2} + \frac{\alpha}{2}. \tag{S38}$$

The embryo-centroid coordinate is therefore

$$r'_{\mathrm{emb},3} = -C_{\mathrm{emb}} \frac{\sin(\alpha/2)}{\alpha/2}, \tag{S39}$$

where

$$C_{\mathrm{emb}} = \frac{a_{\mathrm{emb}}^2 + 4r_{\mathrm{emb}}^2}{4r_{\mathrm{emb}}}. \tag{S40}$$

Because the uniform spherical contribution is centered at the egg origin, the center-of-mass offset of the composite egg is

$$d'_3 = \frac{\Delta m_{\mathrm{oil}} r'_{\mathrm{oil},3} + \Delta m_{\mathrm{emb}} r'_{\mathrm{emb},3}}{m}. \tag{S41}$$

where $r'_{\mathrm{oil},3} = a - a_{\mathrm{oil}}$ from Eq. S24. Its magnitude,

$$d = |d'_3| \tag{S42}$$

is the distance between the geometric center of the egg and its center of mass used in the gravitational-torque calculation.

### Moment of inertia

In the body-fixed principal-axis coordinate system, the mass moment-of-inertia tensor is diagonal. The one-axis model (Eq. S15) therefore requires only the $I_{11}$ component, corresponding to rotation about $x'_1$.

The contribution of the uniform spherical egg, $I_{f,11}$, and the density-contrast contribution of the oil globule, $\Delta I_{\mathrm{oil},11}$, were calculated from the moment of inertia of a sphere, with a parallel-axis correction for the displaced oil globule:

$$I_{f,11} = \frac{2}{5}\rho_f V_{\mathrm{egg}} a^2. \tag{S43}$$

$$\Delta I_{\mathrm{oil},11} = \Delta\rho_{\mathrm{oil}} V_{\mathrm{oil}} \left[\frac{2}{5}a_{\mathrm{oil}}^2 + (a - a_{\mathrm{oil}})^2\right]. \tag{S44}$$

Because the oil globule is represented as a density-contrast contribution relative to the internal fluid, its corresponding moment-of-inertia contribution is denoted $\Delta I_{\mathrm{oil},11}$.

The density-contrast contribution of the embryo was calculated by integrating the $x'_1$ component of the moment of inertia about the geometric center of the egg:

$$\Delta I_{\mathrm{emb},11} = \Delta\rho_{\mathrm{emb}} \int_{V_{\mathrm{emb}}} \left({x'_2}^2 + {x'_3}^2\right) dV. \tag{S45}$$

Using the cylindrical coordinates $r'_1$ and $\phi'_1$ defined in the previous section, the volume element is

$$dV = r'_1\, dr'_1\, dx'_1\, d\phi'_1. \tag{S46}$$

The integration domain was

$$\alpha_1 \le \phi'_1 \le \alpha_2, \qquad -a_{\mathrm{emb}} \le x'_1 \le a_{\mathrm{emb}}, \tag{S47}$$

and

$$r_{\mathrm{emb}} - \sqrt{a_{\mathrm{emb}}^2 - {x'_1}^2} \le r'_1 \le r_{\mathrm{emb}} + \sqrt{a_{\mathrm{emb}}^2 - {x'_1}^2}. \tag{S48}$$

Substituting Eqs. ??, S47, and S48 into Eq. S45 gives

$$\Delta I_{\text{emb},11} = \Delta\rho_{\text{emb}} \int_{\alpha_1}^{\alpha_2} \int_{-a_{\text{emb}}}^{a_{\text{emb}}} \int_{r_{\text{emb}}-\sqrt{a_{\text{emb}}^2 - {x_1'}^2}}^{r_{\text{emb}}+\sqrt{a_{\text{emb}}^2 - {x_1'}^2}} {r_1'}^3 \, dr_1' \, dx_1' \, d\phi_1'. \tag{S49}$$

The integral was evaluated using MATLAB's tiled numerical quadrature, `integral3` with `Method="tiled"`. No fixed spatial grid was prescribed.

The total moment of inertia entering Eq. S15 is therefore

$$I_{11} = I_{f,11} + \Delta I_{\text{oil},11} + \Delta I_{\text{emb},11}. \tag{S50}$$

### Model application and comparison with measurements

#### Stable-orientation solution

A stable-oriented egg does not rotate about its own center in the earthbound frame. Therefore,

$$\omega = 0, \qquad \dot{\omega} = 0. \tag{S51}$$

Consequently, the left-hand side of Euler's equation (Eq. S15) is zero, and the flow-induced torque and the gravitational restoring torque must balance,

$$\tau_{\text{ext},1} + \tau_{\text{int},1} = 0. \tag{S52}$$

Using the torque expressions derived above (Eqs. S17 and S23) and given that $\omega_{rel} = \omega_{cyl}$, this balance gives

$$8\pi\mu a^3 \omega_{\text{cyl}} = mgd \sin\left(\frac{\pi}{2} - \phi_{\text{egg}}\right). \tag{S53}$$

and hence,

$$\phi_{\text{egg}} = \frac{\pi}{2} + \sin^{-1}\left(\frac{8\pi\mu a^3\omega_{\text{cyl}}}{mgd}\right). \tag{S54}$$

A real stable solution exists only when

$$\left|\frac{8\pi\mu a^3\omega_{\text{cyl}}}{mgd}\right| \leq 1. \tag{S55}$$

The limiting cylinder angular velocity for a given egg is therefore

$$\omega_{\text{cyl,crit}} = \frac{mgd}{8\pi\mu a^3}. \tag{S56}$$

Notably, after substituting the neutral buoyancy condition $m = \frac{4}{3}\pi a^3\rho_w$, the critical velocity reduces to

$$\omega_{\text{cyl,crit}} = \frac{\rho_w g d}{6\mu}, \tag{S57}$$

which is independent of egg size and depends only on the center-of-mass offset $d$. Therefore, among neutrally buoyant eggs of the same developmental stage, the transition from stable orientation to revolving motion is determined solely by the degree of bottom-heaviness.

### Testing the stable-orientation model

The stable-orientation model was tested against measurements from 1 DPF eggs classified as stable-oriented. This age group was used because it provided sufficient stable-oriented tracks across the tested cylinder angular velocities. Measurements were grouped by cylinder angular velocity, and the mean egg orientation and standard deviation were calculated for each group. Group means based on fewer than five tracks were excluded from the parameter test but retained in the figure as markers without borders or error bars. Measurements from 2 DPF stable-oriented eggs are shown in Fig. S3A for completeness but were not used to constrain the stable-orientation model.

Because embryo morphology was measured directly, the ranges of $\alpha$ and $a_{\text{emb}}$ used in the parameter test were restricted to the observed ranges of the 76 measurable 1 DPF eggs with partial-torus embryo morphology:

$$107^\circ \le \alpha \le 176^\circ, \tag{S58}$$

$$0.0458\ \text{mm} \le a_{\text{emb}} \le 0.0798\ \text{mm}. \tag{S59}$$

The embryo density was not measured directly and was therefore varied over the prescribed literature-informed range

$$1.03\ \text{mg}\,\text{mm}^{-3} \le \rho_{\text{emb}} \le 1.15\ \text{mg}\,\text{mm}^{-3}. \tag{S60}$$

For each parameter combination, Eqs. S30–S41 were used to calculate the center-of-mass offset, $d$, and Eq. S54 was used to predict the stable egg orientation. The objective function was the unweighted sum of squared angular residuals,

$$\text{SSE} = \sum_{i=1}^{N} \left[\phi_{\text{egg,model}}\left(\omega_{\text{cyl},i}\right) - \overline{\phi}_{\text{egg},i}\right]^2, \tag{S61}$$

where $N$ is the number of cylinder-angular-velocity groups included in the test, $i$ indexes each group, and $\overline{\phi}_{\text{egg},i}$ is the measured mean egg orientation of the $i$-th group.

The stable-angle prediction depends on the internal morphology primarily through the gravitational torque scale $mgd$. Consequently, the stable-orientation measurements do not uniquely identify $\alpha$, $a_{\text{emb}}$, and $\rho_{\text{emb}}$ independently. Within the morphology-constrained parameter space, one parameter combination producing the minimum objective function was

$$\alpha = 154.2^\circ, \qquad a_{\text{emb}} = 0.0573\ \text{mm}, \qquad \rho_{\text{emb}} = 1.150\ \text{mg}\,\text{mm}^{-3}. \tag{S62}$$

This combination produced $mgd = 6.87$ nN mm, an RMSE of 2.21°, and $R^2 = 0.978$.

Both resulting morphology values lie within the measured 1 DPF distributions. The selected value of $\alpha$ was approximately at the 30th empirical percentile, while the selected value of $a_{\mathrm{emb}}$ was approximately at the 36th percentile. The resulting embryo density reached the upper bound of the prescribed range. Because $\rho_{\mathrm{emb}}$ was not measured directly and the individual embryo parameters are not independently identifiable from the stable-angle data, this boundary value should not be interpreted as a direct estimate of embryo density.

The parameter combination in Eq. S62 should therefore be interpreted as one morphology-consistent realization of the gravitational torque scale rather than as a uniquely determined set of embryo properties. Other combinations within the measured morphology ranges can produce the same stable-angle prediction.

### Numerical simulation of revolving eggs

For revolving eggs, Euler's equation (Eq. S13) was integrated numerically in the body-fixed coordinate system. The numerical implementation retained the full vector form of the equation, while the applied torques and initial conditions were restricted to rotation about the $x'_1 = x_1$ axis, normal to the imaging plane. This corresponds to the in-plane rotational motion quantified experimentally.

Using a first-order forward Euler scheme, the angular velocity was updated according to

$$\boldsymbol{\omega}_{n+1} = \mathbf{I}^{-1}\left[\mathbf{I}\boldsymbol{\omega}_n - \Delta t\,\boldsymbol{\omega}_n \times (\mathbf{I}\boldsymbol{\omega}_n) + \Delta t\,(\boldsymbol{\tau}_{\mathrm{int},n} + \boldsymbol{\tau}_{\mathrm{ext},n})\right], \tag{S63}$$

where $n$ denotes the time-step index and $t_n = n\Delta t$.

The angular acceleration over each time step was calculated as

$$\dot{\boldsymbol{\omega}}_{n+1} = \frac{\boldsymbol{\omega}_{n+1} - \boldsymbol{\omega}_n}{\Delta t}, \tag{S64}$$

and the orientation was updated according to

$$\boldsymbol{\theta}_{n+1} = \boldsymbol{\theta}_n + \Delta t\,\boldsymbol{\omega}_{n+1} + \frac{1}{2}\Delta t^2 \dot{\boldsymbol{\omega}}_{n+1}. \tag{S65}$$

Each simulation was initialized with

$$\boldsymbol{\theta}(0) = \mathbf{0}, \qquad \boldsymbol{\omega}(0) = \mathbf{0}. \tag{S66}$$

A time step of $\Delta t = 0.001$ s was used, and each simulation contained 300,000 time steps, corresponding to a simulated duration of 300 s. At each time step, the flow-induced and gravitational restoring torques were recalculated from Eqs. S17 and S23, respectively, using the instantaneous egg angular velocity and orientation.

#### Calculation of revolution frequency and amplitude of relative angular velocity

The first 20% of each simulated time series was discarded to reduce the influence of the initial transient. Because each simulation lasted 300 s, the frequency and amplitude calculations were based on the final 240 s.

The period was obtained from successive local maxima of $\omega_{\text{rel}}(t)$. If $n_k$ denotes the time-step index of the $k$th detected maximum. The mean period was estimated by averaging the time intervals between successive local maxima of $\omega_{\text{rel}}(t)$,

$$\overline{T} = \Delta t \, \frac{1}{N_p - 1} \sum_{k=1}^{N_p - 1} \left( n_{k+1} - n_k \right), \tag{S67}$$

where $N_p$ is the number of detected peaks. The egg revolution frequency was then

$$f_{\text{egg}} = \frac{1}{\overline{T}}. \tag{S68}$$

Simulations with fewer than two detected peaks were classified as non-revolving and assigned $f_{\text{egg}} = 0$.

The simulated amplitude of the relative angular velocity was calculated as follows,

$$A_{\omega,\mathrm{sim}} = \frac{\max(\omega_{\mathrm{rel}}) - \min(\omega_{\mathrm{rel}})}{2}. \quad \text{(S69)}$$

### Parameter-sensitivity simulation sweep

To examine how variation in egg morphology and embryo density affects the predicted rotational dynamics, 100,000 model realizations were generated. For each realization, $\alpha$, $a_{\mathrm{emb}}$, $\rho_{\mathrm{emb}}$, and $\omega_{\mathrm{cyl}}$ were sampled independently from uniform distributions.

Unlike the stable-orientation test, which was restricted to the morphology measured for 1 DPF eggs, the parameter-sensitivity sweep was designed to encompass the morphology observed across both developmental stages. Across measurable 1 DPF and 2 DPF eggs, $\alpha$ ranged from 107.4° to 250.6°, and $a_{\mathrm{emb}}$ ranged from 0.0458 to 0.0970 mm. The simulation bounds were therefore set to the slightly expanded ranges

$$100^\circ \leq \alpha \leq 260^\circ, \quad \text{(S70)}$$

$$0.04\ \mathrm{mm} \leq a_{\mathrm{emb}} \leq 0.10\ \mathrm{mm}. \quad \text{(S71)}$$

The embryo density was sampled over the prescribed literature-informed range

$$1.03\ \mathrm{mg\,mm^{-3}} \leq \rho_{\mathrm{emb}} \leq 1.15\ \mathrm{mg\,mm^{-3}}, \quad \text{(S72)}$$

and the cylinder angular velocity over

$$30\ \mathrm{deg\,s^{-1}} \leq \omega_{\mathrm{cyl}} \leq 150\ \mathrm{deg\,s^{-1}}. \quad \text{(S73)}$$

Because the parameters were sampled independently and uniformly, the resulting combinations represent a sensitivity envelope rather than the observed biological distribution of eggs.

For each realization, the component geometry, internal-fluid density, center-of-mass offset, and moment of inertia were recalculated, after which Euler's equation (Eq. S13) was solved numerically using the procedure described above.

For each simulation, the recorded outputs were the egg orientation $\phi_{\text{egg}}(t)$, egg angular velocity $\omega(t)$, relative angular velocity $\omega_{\text{rel}}(t)$, gravitational restoring torque, and flow-induced torque. For simulations classified as revolving, the egg revolution frequency and relative-angular-velocity amplitude were additionally calculated.

#### Comparison between measured and simulated frequencies and amplitudes

This subsection describes how the simulated revolving egg dynamics were compared with the measurements shown in Fig. 3D for revolution frequency and Fig. 4A for relative angular velocity amplitude.

The frequency comparison used 2 DPF eggs classified as revolving. Measurements from 1 DPF revolving eggs are shown in Fig. S3B for completeness but were not used to calculate the agreement metrics reported for the frequency comparison. Measurements were grouped by cylinder angular velocity, and the corresponding cylinder rotation frequency was calculated as

$$f_{\text{cyl}} = \frac{\omega_{\text{cyl}}}{360^\circ}, \tag{S74}$$

where $\omega_{\text{cyl}}$ is expressed in $\deg\,\text{s}^{-1}$.

For each cylinder-angular-velocity group, the mean and standard deviation of the measured egg revolution frequency, $f_{\text{egg}}$, were calculated. In Fig. 3D, measurement groups containing at least three eggs are shown with error bars and were used to calculate $R^2$ and the mean model bias. Groups containing fewer than three eggs are shown as markers without error bars and were excluded from these calculations.

Simulations with calculated frequencies greater than 0.5 Hz were excluded as numerical peak-detection artifacts. All 107 excluded cases returned exactly $f_{\mathrm{egg}} = 500$ Hz, corresponding to detected maxima separated by two numerical time steps ($\Delta t = 0.001$ s). These traces had negligible relative angular velocity amplitudes ($A_{\omega,\mathrm{sim}} \leq 1.4 \times 10^{-3}$ deg s$^{-1}$, with most values near machine precision), indicating nearly stationary solutions rather than rapidly revolving eggs. Small alternating numerical fluctuations in these traces were identified as successive local maxima by the peak-detection criterion. No simulated frequencies occurred between 0.5 and 500 Hz; the maximum retained frequency was 0.404 Hz. Excluding these artifacts left 57,456 accepted revolving simulations.

The accepted simulations were divided into 20 equally spaced $f_{\mathrm{cyl}}$ bins over 0–0.5 Hz, giving a bin width of 0.025 Hz. For each occupied bin, the mean and standard deviation of $f_{\mathrm{egg}}$ were calculated. The model line in Fig. 3D shows the bin mean, and the shaded region shows the mean plus or minus one standard deviation.

For quantitative comparison, the binned model mean was linearly interpolated to the cylinder frequencies of the measured group means. If $y_i$ is the measured mean frequency, $\widehat{y}_i$ is the corresponding interpolated model frequency and $\overline{y}$ is the mean of the measured frequencies, the coefficient of determination was calculated as

$$R^2 = 1 - \frac{\sum_i (y_i - \widehat{y}_i)^2}{\sum_i (y_i - \overline{y})^2}. \tag{S75}$$

The mean model bias was

$$\mathrm{bias} = \frac{1}{N} \sum_i (\widehat{y}_i - y_i). \tag{S76}$$

where N is the number of cylinder angular velocity groups included in the fit. The model reproduced the measured frequency response with $R^2 = 0.93$ and a mean model-minus-measurement bias of $+0.009$ Hz.

The amplitude required a different measure of agreement because both measured and predicted amplitudes varied over a narrow range. Both $R^2$ and the regression slope are highly sensitive to small point-to-point differences and therefore provided unstable measures of agreement. We therefore quantified model–measurement agreement using RMSE, which directly represents the absolute magnitude of the prediction error in the measured units

$$\mathrm{RMSE} = \sqrt{\frac{1}{N}\sum_i \left(\widehat{y}_i - y_i\right)^2}, \tag{S77}$$

### Estimation of age-dependent center-of-mass offset

The map shown in Fig. 4B describes the predicted magnitude of relative angular velocity as a function of developmental age and cylinder angular velocity, and indicates the boundary between steady-oriented and revolving behavior. Constructing this map requires an estimate of how the center of mass offset, $d$, changes during development. This offset was not measured directly, and the embryo density was not known. We therefore used simulation-based inference (SBI) to estimate an effective embryo density for each experimental observation from its measured morphology and rotational dynamics Cranmer et al. [2020]. The inferred embryo density was then combined with the measured embryo geometry to calculate $d$, and the resulting offsets were regressed against egg age.

SBI is a statistical approach for estimating unknown model parameters by comparing observations with the outputs of numerical simulations. Here, the SBI model was trained using the simulation sweep described above. Each simulation provided a known embryo density and the simulated observation vector produced by that density. The only inferred parameter was the effective embryo density, $\rho_{\mathrm{emb}}$.

The observation vector included two morphological quantities, $\alpha$ and $a_{\mathrm{emb}}$, the imposed cylinder angular velocity, $\omega_{\mathrm{cyl}}$, and two dynamical quantities, $f_{\mathrm{egg}}$ and $A_\omega$:

$$\mathbf{x} = (\alpha,\, a_{\mathrm{emb}},\, \omega_{\mathrm{cyl}},\, f_{\mathrm{egg}},\, A_{\omega})\,. \tag{S78}$$

Age was not included in the observation vector. The age dependence described below was therefore obtained only after estimating the effective embryo density for each observation.

Before training, each component of the observation vector was standardized using the mean and standard deviation of the simulation-sweep dataset; the same transformation was applied later to the experimental observations before inference. Neural posterior estimation was then used to estimate the probability distribution of embryo-density values that are consistent with each observation vector,

$$p\left(\rho_{\mathrm{emb}} \mid \mathbf{x}\right), \tag{S79}$$

where the vertical bar denotes "given". Thus, this distribution describes the possible values of $\rho_{\mathrm{emb}}$ given the measured morphology and rotational dynamics.

Embryo density was inferred using a uniform prior constructed from the simulated density range and extended by 2% on each side, so that all values within the prior were treated as equally likely before conditioning on the observations:

$$\rho_{\mathrm{emb}} \sim \mathcal{U}\,(1.0276, 1.1524)\ \ \mathrm{mg\,mm^{-3}}. \tag{S80}$$

After training, each experimental observation vector was supplied to the trained SBI model, and 500 samples were drawn using Markov chain Monte Carlo from the posterior distribution corresponding to each observation. The observation vector was constructed for revolving eggs only, as stable-oriented eggs do not produce the dynamical quantities $f_{\mathrm{egg}}$ and $A_{\omega}$ required for inference. For each observation $i$, the embryo-density estimate was defined as the posterior mean,

$$\widehat{\rho}_{\mathrm{emb},i} = E[\rho_{\mathrm{emb}} \mid \mathbf{x}_i]\,, \tag{S81}$$

where $\widehat{\rho}_{\mathrm{emb},i}$ denotes the estimated effective embryo density for observation $i$, $\mathbf{x}_i$ is the corresponding observation vector, and $E[\cdot]$ denotes the expectation, or probability-weighted mean, of the posterior distribution. In practice, this expectation was approximated by the arithmetic mean of the 500 posterior samples drawn for each observation.

The posterior mean density estimate was then combined with the measured values of $\alpha_i$ and $a_{\mathrm{emb},i}$ in the geometrical egg model described above. This produced one estimate of the center-of-mass offset, $d_i$, for each experimental observation. Across the 49 observations of revolving eggs, the estimated offsets ranged from approximately $4.02 \times 10^{-4}$ to $1.27 \times 10^{-3}$ mm.

The estimated offsets were then paired with the recorded age of each egg, $t_i$, in hours post-fertilization. An ordinary least-squares line was fitted to all 49 pairs $(t_i, d_i)$,

$$d(t) = -1.388 \times 10^{-5} t + 1.218 \times 10^{-3} \text{ mm}, \tag{S82}$$

where $t$ is in hours post-fertilization. Equation S82 therefore describes the developmental trend in center of mass offset estimated from the combination of SBI-inferred embryo density and measured embryo geometry.

### Construction of the age-rotation map

The age-dependent relation $d(t)$ was inserted into the torque-balance criterion (Eq. S55) to construct Fig. 4B. For a given egg, the condition presented in Eq. S55 was used to determine the critical $\omega_{cyl}$ above which the egg revolves and under which the egg maintains steady orientation. This condition defines the boundary shown in Fig. 4B:

$$\omega^*_{\mathrm{cyl}}(t) = \frac{mgd(t)}{8\pi\mu a^3}. \tag{S83}$$

For cases with $\omega_{cyl} < \omega^*_{\mathrm{cyl}}(t)$, the egg relative angular velocity magnitude is $|\omega_{rel}| = |\omega_{cyl}|$. For $\omega_{\mathrm{cyl}} > \omega^*_{\mathrm{cyl}}(t)$, the plotted value represents the maximum magnitude of the relative angular velocity predicted by the torque balance.

At the point where $|\omega_{\mathrm{rel}}|$ reaches its maximum, the angular acceleration term in Eq. S13 is zero. The rotational equation therefore reduces locally to a balance between the flow induced and gravitational torques,

$$8\pi\mu a^3\omega_{\mathrm{rel}} + mgd(t)\sin\left(\frac{\pi}{2} - \phi_{\mathrm{egg}}\right) = 0. \tag{S84}$$

The magnitude of the restoring torque is maximal when

$$\left|\sin\left(\frac{\pi}{2} - \phi_{\mathrm{egg}}\right)\right| = 1. \tag{S85}$$

Thus, the maximum magnitude of the relative angular velocity in the revolving regime is

$$|\omega_{\mathrm{rel}}|_{\mathrm{max}} = \frac{mgd(t)}{8\pi\mu a^3} \tag{S86}$$

This value is equal to the critical cylinder angular velocity in Eq. S83, because both are set by the maximum gravitational restoring torque. Combining the two regimes, the relative angular velocity plotted in Fig. 4B was therefore

$$\frac{|\omega_{\mathrm{rel}}|\,(t, \omega_{\mathrm{cyl}}) = \begin{cases} \omega_{\mathrm{cyl}}, & \omega_{\mathrm{cyl}} \leq \omega^*_{\mathrm{cyl}}(t), \\ \mathrm{mgd(t)} & \end{cases}}{8\pi\mu a^3}$$

$\omega_{\mathrm{cyl}} > \omega^*_{\mathrm{cyl}}(t)$. (S87)

Eq. 0.1 was evaluated on a regular grid spanning ages of 10–50 hpf and cylinder angular velocities of 30–150 deg $\mathrm{s}^{-1}$. The resulting field of $|\omega_{\mathrm{rel}}|$ was shown as a filled contour map, and the equality condition in Eq. S55 was plotted as the boundary between stable-oriented and revolving behavior.

Table S1: Model parameters and quantities used in the rotational-dynamics model.

| Symbol | Description | Value or definition |
|---|---|---|
| $a$ | Egg radius | 0.5 mm |
| $a_{\text{oil}}$ | Oil globule radius | 0.1 mm |
| $a_{\text{emb}}$ | Embryo minor radius | Measured/model parameter |
| $r_{\text{emb}}$ | Embryo major radius | $a - a_{\text{emb}}$ |
| $\alpha$ | Angular extent of partial torus embryo | Measured/model parameter |
| $\alpha_1, \alpha_2$ | Angular limits of embryo segment | $\alpha_2 - \alpha_1 = \alpha$ |
| $\rho_w$ | Seawater density | 1.03 $\text{mg}\,\text{mm}^{-3}$ |
| $\rho_{\text{oil}}$ | Oil globule density | 0.90 $\text{mg}\,\text{mm}^{-3}$ |
| $\rho_{\text{emb}}$ | Embryo density | Model parameter |
| $\rho_f$ | Density of remaining internal fluid | Calculated from neutral buoyancy |
| $\Delta\rho_{\text{oil}}$ | Oil globule density contrast | $\rho_{\text{oil}} - \rho_f$ |
| $\Delta\rho_{\text{emb}}$ | Embryo density contrast | $\rho_{\text{emb}} - \rho_f$ |
| $V_{\text{egg}}$ | Egg volume | $\frac{4}{3}\pi a^3$ |
| $V_{\text{oil}}$ | Oil globule volume | $\frac{4}{3}\pi a_{\text{oil}}^3$ |
| $V_{\text{emb}}$ | Embryo volume | $2\pi^2 r_{\text{emb}} a_{\text{emb}}^2 \frac{\lvert\alpha\rvert}{2\pi}$ |
| $m$ | Total egg mass | $\rho_w V_{\text{egg}}$ |
| $\Delta m_{\text{oil}}$ | Oil density-contrast mass contribution | $V_{\text{oil}}\Delta\rho_{\text{oil}}$ |
| $\Delta m_{\text{emb}}$ | Embryo density-contrast mass contribution | $V_{\text{emb}}\Delta\rho_{\text{emb}}$ |
| $\mathbf{r}'_{\text{oil}}$ | Oil globule centroid position | $[0, 0, a - a_{\text{oil}}]^{\mathsf{T}}$ |
| $\mathbf{r}'_{\text{emb}}$ | Embryo centroid position | Calculated from partial-torus geometry |
| $d'_3$ | Center of mass coordinate along $x'_3$ | Calculated from density contrast contri[illegible] |
| $d$ | Center of mass offset magnitude | $d = \lvert d'_3\rvert$ |
| $I_{11}$ | Moment of inertia about $x'_1$ | Calculated from component geometry a[illegible] |
| $\mu$ | Dynamic viscosity | 1.09 $\text{nN}\,\text{s}\,\text{mm}^{-2}$ |
| $\nu$ | Kinematic viscosity | $\mu/\rho_w$ |
| $g$ | Gravitational acceleration | $0.98 \times 10^4$ $\text{mm}\,\text{s}^{-2}$ |
| $\omega_{\text{cyl}}$ | Cylinder angular velocity about $X_1$ | Relative to laboratory frame; experimen[illegible] |
| $\omega$ | Egg angular velocity about $x'_1 = x_1$ | Relative to laboratory frame |
| $\omega_{\text{rel}}$ | Water angular velocity relative to egg about $x'_1 = x_1$ | $\omega_{\text{cyl}} - \omega$ |
| $\phi_{\text{egg}}$ | Egg orientation angle | Angle between $+x_2$ and $+x'_3$ |

Table S2: Agreement between model predictions and experimental measurements.

| Analysis | $R^2$ | Regression slope | RMSE | Normalized RMSE |
|---|---|---|---|---|
| Stable orientation | 0.97 | 0.95 | 2.53 deg | 2.02% |
| Rotation frequency | 0.93 | 0.78 | 0.019 Hz | 7.89% |
| Angular-velocity amplitude | * | * | $4.3928^\circ\,\text{s}^{-1}$ | 7.26% |

Normalized RMSE was calculated relative to the mean measured value. *For the amplitude analysis, both the regression slope and $R^2$ are highly sensitive to small point-to-point differences because the measured and predicted amplitudes span restricted ranges; in particular, the model predictions span only 0.27% of their mean. They are therefore not reported, and RMSE and normalized RMSE are used as the primary measures of model–measurement agreement.

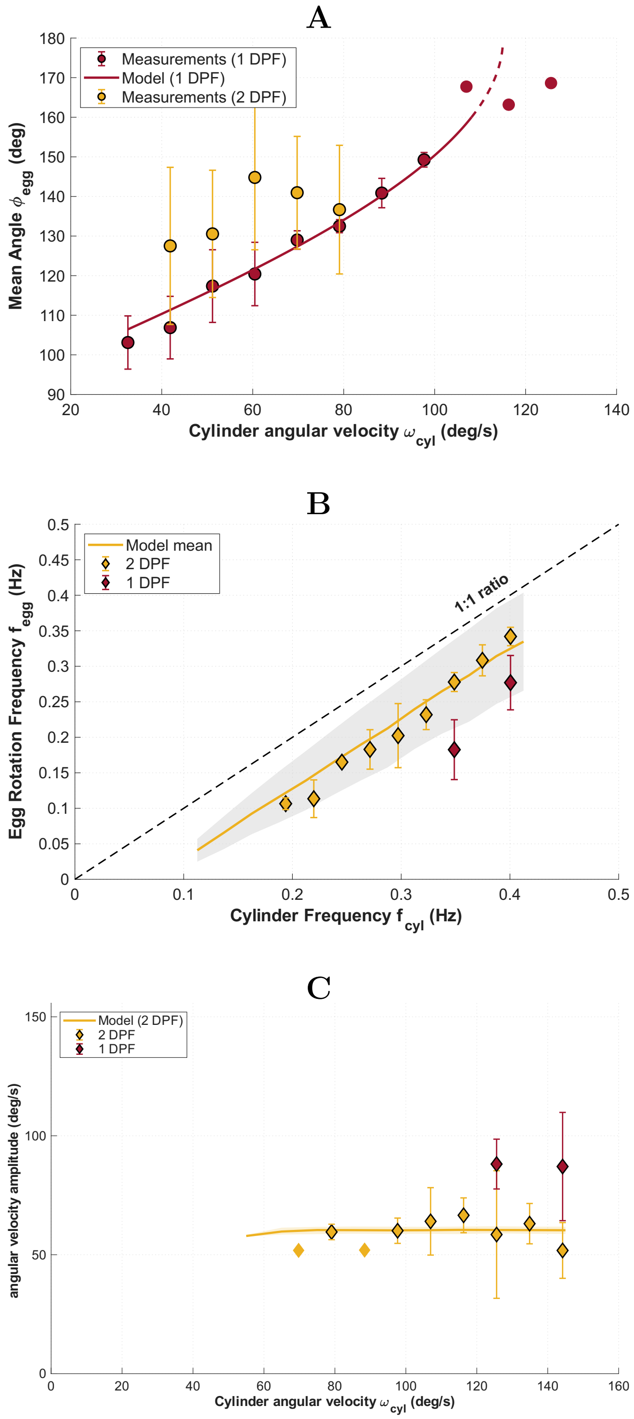


Figure S3: Comparison between measured and modeled rotational dynamics for both developmental age groups. (A) Stable egg orientation as a function of cylinder angular velocity. (B) Egg revolution frequency as a function of cylinder rotation frequency. (C) Relative-angular-velocity amplitude as a function of cylinder angular velocity.

### Supplementary Movies

**Movie S1.** Type legend for the movie here.

**Movie S2.** Type legend for the other movie here.

### Supplementary Datasets

**Dataset S1** (`dataset_one.txt`): Type or paste legend here.

**Dataset S2** (`dataset_two.txt`): Type or paste legend here.

Talmo D. Pereira, Nathaniel Tabris, Arie Matsliah, David M. Turner, Junyu Li, Shruthi Ravindranath, Eleni S. Papadoyannis, Edna Normand, David S. Deutsch, Z. Yan Wang, Grace C. McKenzie-Smith, Catalin C. Mitelut, Marielisa Diez Castro, John D'Uva, Mikhail Kislin, Dan H. Sanes, Sarah D. Kocher, Samuel S.-H. Wang, Annegret L. Falkner, Joshua W. Shaevitz, and Mala Murthy. SLEAP: A deep learning system for multi-animal pose tracking. *Nature Methods*, 19(4):486–495,

April 2022. ISSN 1548-7091, 1548-7105. doi: 10.1038/s41592-022-01426-1. URL https://www.nature.com/articles/s41592-022-01426-1.

E.R. Davies. Circle Detection. In *Machine Vision*, pages 283–313. Elsevier, 2005. ISBN 978-0-12-206093-9. doi: 10.1016/B978-012206093-9/50013-7. URL https://linkinghub.elsevier.com/retrieve/pii/B9780122060939500137.

I. Rønnestad, W. M. Koven, A. Tandler, M. Harel, and H. J. Fyhn. Energy metabolism during development of eggs and larvae of gilthead sea bream (Sparus aurata). *Marine Biology*, 120(2):187–196, September 1994. ISSN 0025-3162, 1432-1793. doi: 10.1007/BF00349678. URL http://link.springer.com/10.1007/BF00349678.

H. Okan Kamaci, Sahin Saka, and Kursat Firat. The Cleavage and Embryonic Phase of Gilthead Sea Bream (Sparus aurata Linnaeus, 1758) Eggs. 22(1):205–209, 2005. ISSN 1300 - 1590.

Kyle Cranmer, Johann Brehmer, and Gilles Louppe. The frontier of simulation-based inference. *Proceedings of the National Academy of Sciences*, 117(48):30055–30062, December 2020. ISSN 0027-8424, 1091-6490. doi: 10.1073/pnas.1912789117. URL https://pnas.org/doi/full/10.1073/pnas.1912789117.
